# One gene, multiple ecological strategies: a biofilm regulator is a capacitor for sustainable diversity

**DOI:** 10.1101/2020.05.02.074534

**Authors:** Eisha Mhatre, Daniel J. Snyder, Emily Sileo, Caroline B. Turner, Sean W. Buskirk, Nico L. Fernandez, Matthew B. Neiditch, Christopher M. Waters, Vaughn S. Cooper

## Abstract

Many bacteria cycle between sessile and motile forms in which they must sense and respond to internal and external signals to coordinate appropriate physiology. Maintaining fitness requires genetic networks that have been honed in variable environments to integrate these signals. The identity of the major regulators and how their control mechanisms evolved remain largely unknown in most organisms. During four different evolution experiments with the opportunist betaproteobacterium *Burkholderia cenocepacia* in a biofilm model, mutations were most frequently selected in the conserved gene *rpfR*. RpfR uniquely integrates two major signaling systems -- quorum sensing and the motile-sessile switch mediated by cyclic-d-GMP -- by two domains that sense, respond to, and control synthesis of the autoinducer cis-2-dodecenoic acid (BDSF). The BDSF response in turn regulates activity of diguanylate cyclase and phosphodiesterase domains acting on cyclic-di-GMP. Parallel adaptive substitutions evolved in each of these domains to produce unique life history strategies by regulating cyclic-di-GMP levels, global transcriptional responses, biofilm production, and polysaccharide composition. These phenotypes translated into distinct ecology and biofilm structures that enabled mutants to coexist and produce more biomass than expected from their constituents grown alone. This study shows that when bacterial populations are selected in environments challenging the limits of their plasticity, the evolved mutations not only alter genes at the nexus of signaling networks but also reveal the scope of their regulatory functions.

**Significance statement:** Many organisms including bacteria live in fluctuating environments requiring attachment and dispersal. These lifestyle decisions require multiple external signals to be processed by several genetic pathways, but how they are integrated is largely unknown. We conducted multiple evolution experiments totaling >20,000 generations with *Burkholderia cenocepacia* populations grown in a model of the biofilm life cycle and identified parallel mutations in one gene, *rpfR*, that is a conserved central regulator. Because RpfR has multiple sensor and catalytic domains, different mutations can produce different ecological strategies that can coexist and even increase net growth. This study demonstrates that a single gene may coordinate complex life histories in biofilm-dwelling bacteria and that selection in defined environments can reshape niche breadth by single mutations.

## Introduction

Bacteria have experienced strong selection over billions of generations to efficiently and reversibly switch from free-swimming to surface-bound life. The record of this selection is etched in the genomes of thousands of species, many of which have tens or even hundreds of genes that govern this lifestyle switch (1). At the nexus of this switch in the majority of bacteria is the second messenger molecule cyclic diguanylate monophosphate (c-di-GMP). Many genes synthesize, degrade, or directly bind and respond to c-di-GMP that in high concentrations promotes a sessile lifestyle and biofilm production and in low concentrations promotes a solitary, motile life. Those genomes with the greatest apparent redundancy in this signaling network demonstrate the highest plasticity along this motile-sessile axis (2). For instance, in *Vibrio cholerae*, there are 41 distinct diguanylate cyclases (DGCs) that synthesize c-di-GMP and 31 different phosphodiesterases (PDEs) that degrade this molecule (1).

Recent theory and experiments suggest that the evolution of this apparent redundancy is driven by the need to integrate many signal inputs generated in fluctuating environments and also produce appropriate outputs in response (3). However, the question remains how so many enzymes that produce or degrade c-di-GMP can be maintained with distinct roles. One explanation is that some DGCs or PDEs exert a dominant effect in certain environmental conditions over the rest of the network. A screen of a complete set of gene knockouts in a low-temperature environment found that only six DGCs were primary contributors to increased levels of c-di-GMP in *V. cholerae* (4, 5). Similar approaches in *Pseudomonas*, which generally contain 40 or more genes encoding DGC, PDE, or both domains, suggest that these enzymes form complexes that are tailored to the prevailing sensed condition (6, 7). An active frontier in this field now seeks to define and characterize the external cues that activate these specific regulatory circuits, that is, how does the single second messenger c-di-GMP function as the decisive node for variable bacterial life-history strategies based on cues originating outside the cell?

Both of these questions – what gene products dominate in c-di-GMP signaling, and how do they integrate external signals – motivate this study of evolved populations of *B. cenocepacia*, an opportunistic and metabolically versatile betaproteobacterium that is especially threatening to persons with cystic fibrosis (8). Evidence is mounting that a few of the 25 potential DGCs or PDEs in *B. cenocepacia* are central to this network (9). An early mutant screen of *B. cenocepacia* genes identified one gene, *yciR*, as one of several that increased biofilm production (10). This gene was later renamed *rpfR* (regulator of pathogenicity factors) based on its homolog in *Xanthomonas campestris* and was shown to have both DGC and PDE domains (11). Importantly, this study also identified a PAS sensor domain in RpfR that binds the autoinducer molecule cis-2-dodecenoic acid, otherwise known as *Burkholderia* diffusible signal factor (BDSF) (11). Most recently, a study of deletion mutants of all putative DGC and PDE proteins in *B. cenocepacia* str. J2135 pointed to *rpfR* as being of particular importance (9). RpfR is now recognized as a bifunctional protein consisting of both DGC and PDE domains as well as two sensor domains, the second of which we recently discovered (12). One sensor is a Per-Arnt-Sim (PAS) domain that binds BDSF (11) which then stimulates the PDE domain that cleaves c-di-GMP to pGpG and GMP (Fig. 1A). Thus, BDSF, like other DSFs, promotes biofilm dispersal by decreasing cellular c-di-GMP levels.

**Fig 1.**
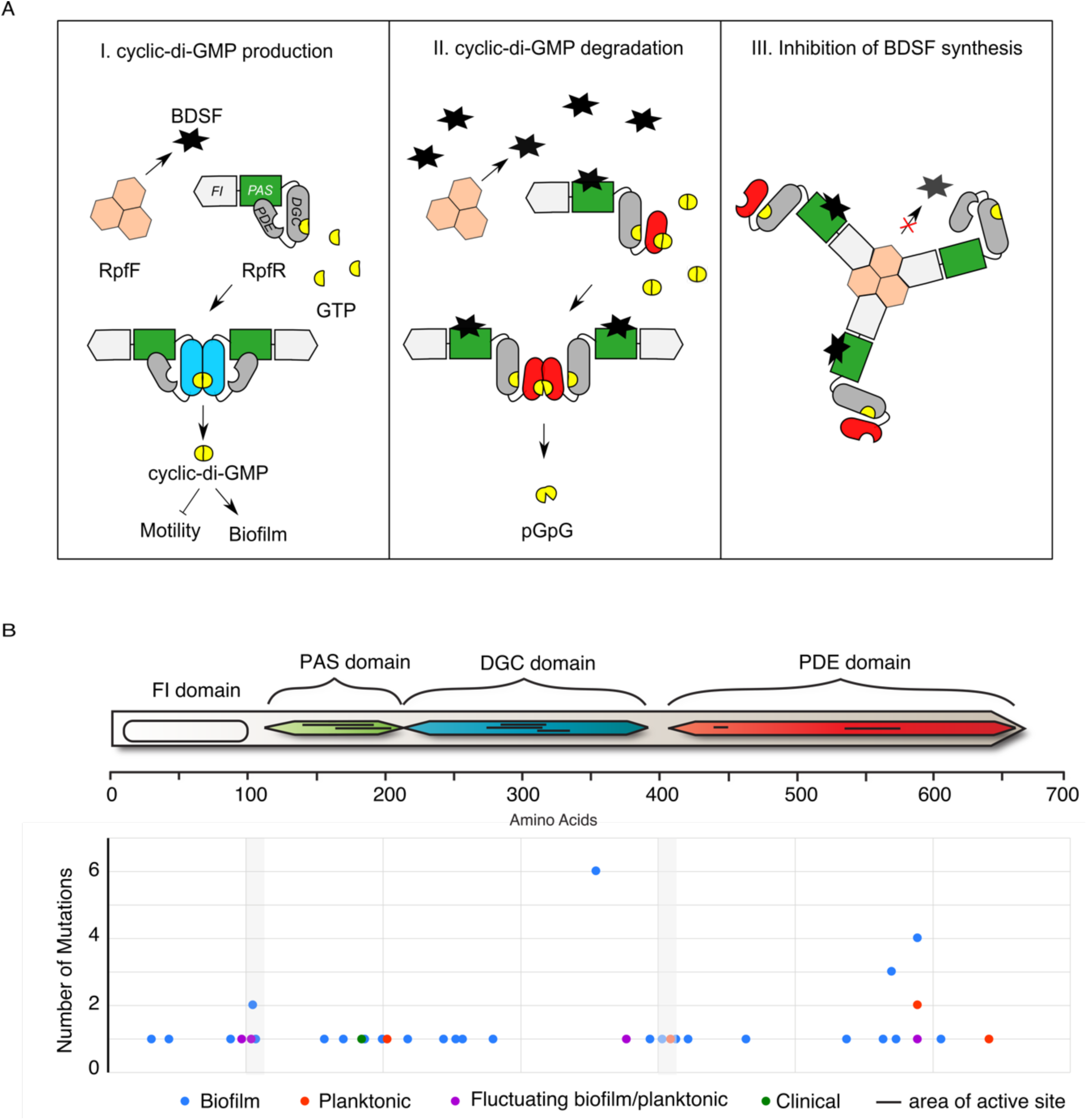
RpfR is the dominant target of selection in *Burkholderia* biofilms. (A) Hypothesized model of BDSF signaling and c-di-GMP metabolism by the RpfR-RpfF regulon. RpfR consists of four domains: i) RpfF-inhibiting or FI domain, ii) Per-Arnt-Sim (PAS) sensor, iii) a diguanylate cyclase (DGC) domain with GGDEF motif and iv) a phosphodiesterase (PDE) domain with EAL motif. Its adjacent gene product RpfF is an enoyl-CoA-hydratase that produces *Burkholderia* Diffusible Signal Factor (BDSF). Panel I: The DGC domain, when active (blue), synthesizes c-di-GMP, a second messenger that regulates biofilm formation and motility. Panel II: BDSF binds the PAS domain and induces c-di-GMP degradation by activating the PDE domain (red). Panel III: The FI domain of RpfR binds RpfF and forms a complex that inhibits BDSF production (12, 14, 47) (B) Evolved *rpfR* mutations during experimental selection in biofilm (blue), planktonic (red), alternating biofilm and planktonic growth (purple) and chronic infections of the CF lung (green). Mutations are disproportionately enriched in linker regions (gray shading) between the sensor and catalytic domains, and at four residues (Table 1).

Discovery of the second sensory domain was partly informed by our evolution experiments with *B. cenocepacia* in our biofilm bead model, in which bacteria are selected to colonize a polystyrene bead that is transferred each day to a new test tube containing media and a fresh bead (13). Evolved *rpfR* mutants from these studies led us to identify an additional N-terminal domain of this protein that was previously uncharacterized in the protein database (12). We named this domain the RpfF-Inhibitory domain, or FI domain, because it binds RpfF, the thioesterase that produces BDSF that is encoded by the adjacent gene. When RpfR-FI binds RpfF it negatively regulates its production of BDSF (Fig. 1A) (12). This finding led us to hypothesize that *rpfR* was a focus of selection not only because it governs c-di-GMP-mediated biological processes but also BDSF-related quorum sensing (11, 14).

**Table 1:** Distribution of *rpfR* mutations by domains and statistical enrichment in linker regions (X^2^ = 10.47, df=1, p = 0.0012). Mutation probability is calculated for 51 SNPs only from Table S1, assuming probability proportionate to domain size. A full list of mutations is present in Table S1.

| Domain | Observed mutations | Expected mutations | Residues |
| --- | --- | --- | --- |
| FI | 3 | 7.125 | 1-95 |
| PAS | 8 | 8.25 | 113-222 |
| GGDEF (DGC) | 13<sup>a</sup> | 12.225 | 235-397 |
| Linker regions | 8 | 2.175 | 96-113,<br>222-235<br>398-409 |
| EAL (PDE) | 19<sup>b</sup> | 19.2 | 409-664 |
| GGDEF+EAL | 1 | n/a | 235-664 |
| Full deletion + rpfF | 18 | n/a |  |
| Full deletion | 2 | n/a | 1-680 |
| Sum, SNPs only | 51 |  |  |
| Sum | 72 |  |  |

This integration of multiple regulatory roles within one gene raises an important evolutionary question: how does natural selection coordinate the functions of its protein domains given their biochemical opposition (synthesize or degrade c-di-GMP) and their capacity to produce different life histories (stick or swim)? Addressing this question is experimentally intractable by conventional methods using knockout or deletion mutations because they usually obscure effects of individual protein domains and cannot address how altered residues of a broadly conserved gene like *rpfR* influence specific function (15, 16). The point mutations in different RpfR domains that evolved during our long-term evolution experiment encode more nuanced information. Not only did these mutants increase fitness in a model of the biofilm lifestyle, they also coexisted for hundreds of generations, suggesting they produced different phenotypes that did not compete for the same niche (17, 18). Further, in a separate study we discovered a *rpfR* mutation that associated with increased biofilm production and genetic diversification during a 20-year chronic *B. multivorans* infection of a cystic fibrosis patient (19). In both scenarios, the *rpfR* mutations cooccurred with other mutations along their evolutionary trajectories, leaving their independent contributions to fitness and gene function yet to be determined. Here, we use a combination of directed genetics, transcriptomics, and assays of microbial ecology, physiology, and fitness in multiple environments to understand how *rpfR* functions as a regulatory node and why mutations in this system predictably evolve in our biofilm model and perhaps also during infections.

## Results

### Unprecedented parallel selection for *rpfR* mutations during evolution experiments

From previous evolution experiments (13, 17, 20, 21) with *B. cenocepacia* grown in our bead model that simulates the biofilm life cycle, we identified at least 72 *rpfR* mutations in 32 independent populations that affected multiple protein domains (Fig. 1B and Table S1). Mutations in *rpfR* were always among the first mutations to rise to high frequency (>25%) in each experiment and were associated with increased competitive fitness and biofilm production (17, 18). The mutation spectrum demonstrates strong selection for altered or eliminated protein function: 45/46 nucleotide substitutions were nonsynonymous and 26 were deletion mutations or premature stop codons. All but two deletions removed both *rpfR* and the adjacent *rpfF* gene, suggesting that selection acted upon interactions between these two gene products. The distribution of SNPs was also non-random and significantly enriched in linker regions between the four domains rather than in the catalytic or sensory sites themselves (X^2^ = 10.47, df=1, p = 0.0012, Table 1 and Fig. 1B). This result suggests selection for altered interactions between functional domains rather than for disrupting BDSF sensing or c-di-GMP catalysis. Among the 13 mutations in the DGC domain, 8 occurred at Y355 or R377, pointing to the functional importance of these residues. Further, 10 mutations affecting the phosphodiesterase EAL domain occurred in just two positions, S570 (3) and F589 (7). In total, selection acted on the *rpfR* sequence with remarkable precision that prompted further study of their functional roles.

In the long-term evolution experiment, three *rpfR* mutants arose in the same population, coexisted during long-term biofilm selection, and associated with different ecology, which suggested that these mutations were not functionally equivalent (13). These mutants, A106P in the region linking the FI and PAS domains, Y355D in the DGC domain, and a deletion mutant of both *rpfR* and *rpfF* (or a functionally equivalent *de novo* evolved mutant) also evolved in parallel among replicate populations and became a major focus of this study. Together, these findings suggested that selection could produce multiple, discrete phenotypes by altering different domains of a dominant c-di-GMP regulator.

### Biofilm and c-di-GMP levels vary with mutated *rpfR* domains

We introduced the evolved point mutations or targeted deletions into the ancestral HI2424 strain (Table S2 and S3) and confirmed that they were otherwise isogenic by whole-genome sequencing. Hereafter, we refer to these engineered genotypes as evolved mutants. Further, to test the contributions of each sensor and enzymatic domain, we constructed deletions of the FI domain (1-95aa) and alanine replacements predicted to eliminate diguanylate cyclase activity (GGDAF, equivalent to E319A), or phosphodiesterase activity (AAL or E443A). We also deleted *rpfR* in its entirety, the adjacent BDSF synthase *rpfF*, or both these genes. Because a 95-gene deletion removing both *rpfR* and *rpfF* repeatedly evolved during our experiments and was available before successful construction of the *ΔrpfFR* genotype, some experiments were conducted with this *ΔrpfRF*+93 genotype (Table S2). We subsequently competed these two genotypes and found their fitness to be statistically indistinguishable (t = 0.38, df = 10, p = 0.12, Supplementary Data).

A handy screen for elevated c-di-GMP is rugose colony morphology or increased uptake of Congo Red dye, both of which result from the increased polysaccharide production often associated with high c-di-GMP (22). The evolved point mutants (A106P and Y355D) displayed increased uptake of Congo Red dye on morphology plates (Fig 2A) and all evolved colonies produced a characteristic studded center and smooth periphery in contrast with the smooth phenotype of WT (Fig. S1A). These colony phenotypes correlated with increased biofilm production and reduced motility (Fig. 2B and S1B). Similar phenotypes were observed in the engineered AAL and *ΔrpfF* mutants (Fig. S1), which should eliminate the PDE domain activity and BDSF production that activates the PDE domain, respectively, increasing c-di-GMP levels. To test these predictions, we quantified *in vivo* levels of intracellular c-di-GMP at both 12 and 24h from planktonic and biofilm cultures of each mutant (Fig. 2CD and Table S4). First, we learned that absolute values of the signal were generally greater at 24h in denser biofilms, but relative differences (values divided by WT value) were greater at 12h when colonization of the plastic beads accelerates in our model (17, 23). Second, the A106P mutant of the FI-PAS linker region produces modest but consistent increases in c-di-GMP across conditions, suggesting this mutant interferes with PAS-mediated activation of the PDE. Third, as predicted, the AAL mutant that should disable the PDE domain and the *ΔrpfF* mutant that produces no BDSF to activate the PDE domain both increases c-di-GMP. Fourth, the evolved *ΔrpfRF*+93 mutant produced elevated c-di-GMP in biofilms at 12 h and in planktonic cultures at 24h, which suggests that losing the PDE activity of RpfR unmasks contributions of other DGCs. Interestingly, deleting only *rpfR* did not significantly alter c-di-GMP levels in biofilms but did increase levels in planktonic cultures, suggesting that functional RpfF in the absence of RpfR affects the c-di-GMP pool in an unknown manner. Finally, the evolved Y355D mutant of the DGC domain produced the highest levels of c-di-GMP, suggesting this is a gain-of-function mutation in a domain thought to be nonfunctional (24, 25). To test this prediction, we constructed a GGDAF mutation that should disable the DGC domain in the Y355D mutant (Y355D-GGDAF) and found, as expected, it produced WT levels of c-di-GMP (Fig. S2A). This result demonstrated that the RpfR DGC domain is directly responsible for the high c-di-GMP levels in Y355D. Together, these results indicate that evolved genotypes produce different basal levels of c-di-GMP depending on the affected domain and alter production depending on their environment. In broader terms, growth of *B. cenocepacia* in biofilms can select for differentiated biofilm-associated phenotypes caused by single *rpfR* mutations.

**Fig. 2.**
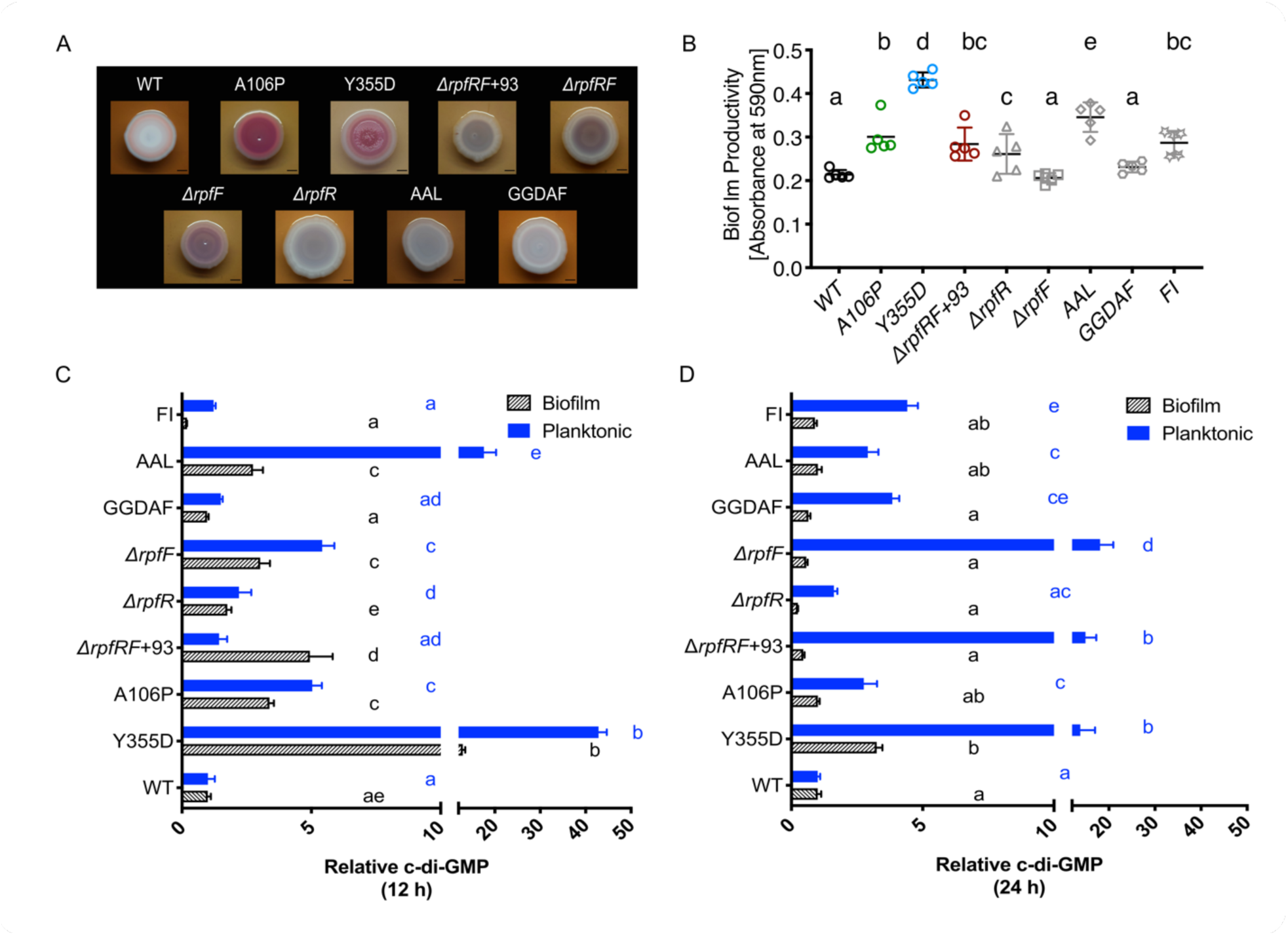
Evolved and engineered *rpfR* genotypes produce diverse c-di-GMP-regulated phenotypes. (A) Colony characteristics of evolved and engineered mutants on Congo Red, tryptone agar plates. (B) Biofilm productivity measured by crystal violet staining. Evolved mutants are in colors and engineered mutants are in grey. (C) Relative levels of c-di-GMP to WT measured at 12 (C) and (D) 24 hours in biofilm (black letters) and planktonic conditions (blue letters). Error bars are 95% c.i. Letters denote significant differences between mutants (one-way ANOVA, post hoc comparisons q value< 0.05).

### Fitness in the biofilm model relates to c-di-GMP levels

We predicted that varied c-di-GMP levels and associated differences in biofilm matrix production contributed to fitness. Evolved and engineered mutants were competed against the WT strain in equal ratios and demonstrated significant variation in fitness, with the Y355D mutant the most fit (Fig. 3). Overall, fitness in the biofilm model at 24h, when development matures, positively correlated with c-di-GMP levels at 12h, when rates of attachment accelerate (Fig. 3A). However, the rate of fitness increases decelerate with increasing c-di-GMP levels, especially among evolved mutants, suggesting diminishing returns (Figure 3A). The A106P mutant was disproportionately more fit at 24h, implying additional advantages of this genotype affecting the FI-PAS linker region, yet fitness of *ΔrpfF* was equivalent to WT in biofilm despite very high c-di-GMP levels (Fig. 3B). Further, the evolved *ΔrpfRF*+93 and the engineered *ΔrpfR* genotypes were more fit against WT despite modest increases in c-di-GMP. Many of the mutants were also more fit against the WT under planktonic growth conditions, which is a necessary component of our bead model that requires dispersal, but fitness benefits were lower and less variable among mutants than those in biofilm conditions (Fig. S3). The loss of PDE activity (AAL) increased c-di-GMP levels, as predicted, and also greatly increased fitness (Fig. 3AB). This strong benefit suggests that the dominant role of RpfR is its PDE activity, as has been shown in orthologs of other species (3, 26)

**Fig. 3.**
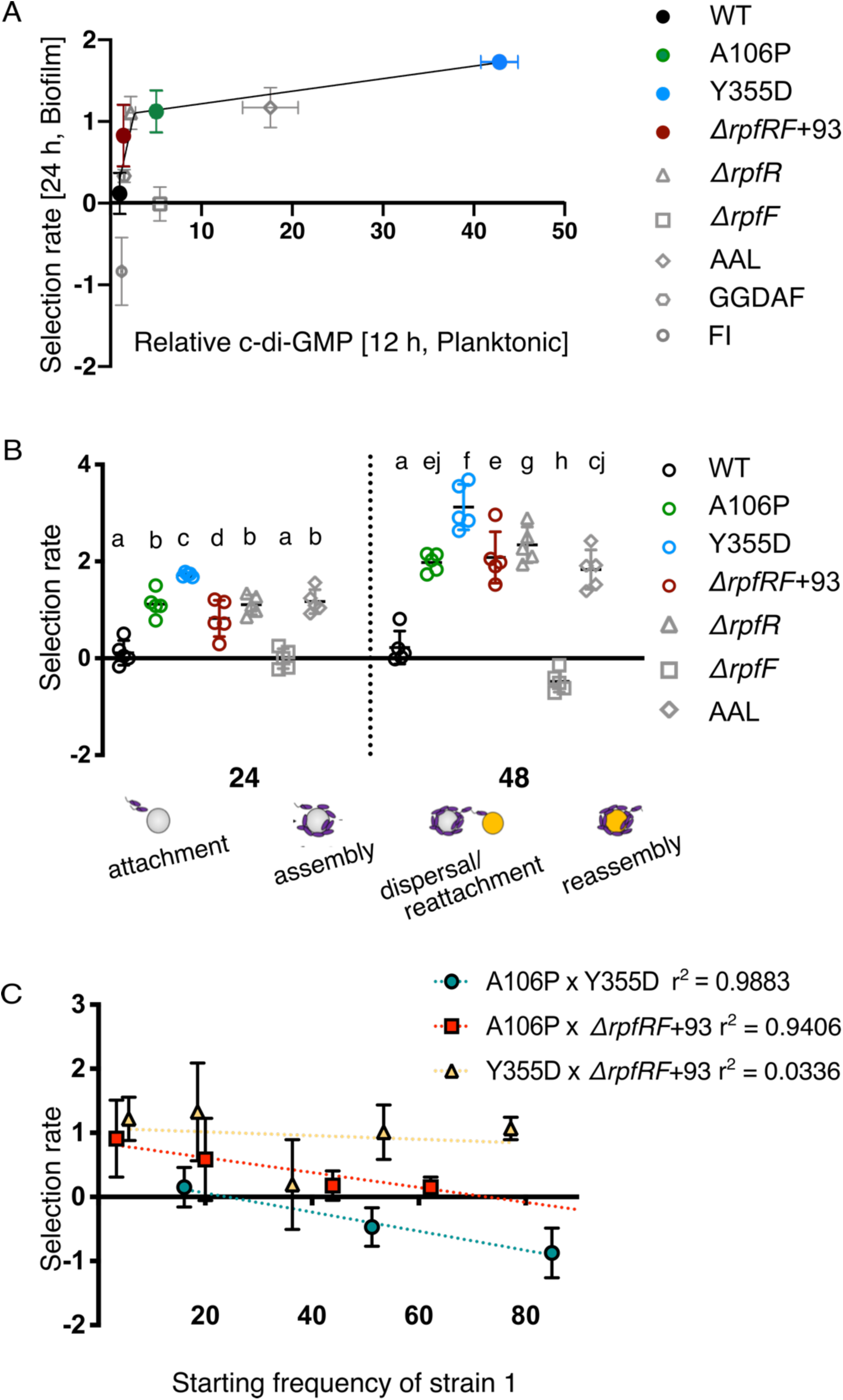
Fitness of *rpfR* genotypes as a function of c-di-GMP levels. (A) Non-linear relationship (segmental linear regression, r^2^ = 0.77) between c-di-GMP production at 12 h and fitness in biofilm at 24 h for evolved mutants. Engineered mutants shown in grey, not included in function. (B) Relative fitness vs. WT during 24h and 48 h in the biofilm model. Different letters indicate significant differences between genotypes by posthoc testing following ANOVA. (C) Fitness differences between *rpfR* mutants after 24h of competition starting from different starting frequencies. The intersection between the regression lines and the x-axis is the predicted mutant frequencies at equilibrium. Error bars are 95% c.i.

### Biofilm ecology: (i) coexistence of *rpfR* mutants

The sustained coexistence of different *rpfR* mutants in evolving biofilm populations (18) could be explained by niche differentiation within the biofilm life cycle. If these niches support populations of different sizes, the fitness of different genotypes should depend on their relative frequencies and be able to invade one another when rare, also known as negative-frequency-dependent selection (NFDS) (27). We tested this hypothesis by competing each evolved genotype versus the others after 24 hours in the biofilm and found support for this model (Fig. 3C). Both A106P and Y355D can invade one another when introduced at low frequency, with a predicted equilibrium frequency of 1:4 A106P: Y355D (Fig. 3C, linear regression analysis y = −0.0149*x + 0.3599, r^2^= 0.9883). Further, the A106P and *ΔrpfRF*+93 or *ΔrpfR* mutants show comparable high fitness in competition with WT (Fig. 2A) but may coexist via NFDS when co-cultured (Fig. 3C, linear regression y = −0.01164*x + 0.8477, r^2^= 0.9406 and Supplementary fig. 4B, y = −0.02265*x + 0.945, r^2^= 0.6421). However, the Y355D mutant was significantly more fit than the *ΔrpfRF*+93 genotype that ultimately displaced it during the long-term evolution experiment (Fig. 3C yellow, linear regression y = −0.0029*x + 1.074, r^2^= 0.0336). High Y355D fitness is consistent with its sweep to high frequency (13) and parallel evolution (Table 1), but this cannot explain why *ΔrpfRF*+93 ultimately displaced Y355D. Prior studies indicated that the spread of other mutations within the *ΔrpfRF*+93 lineage increased its relative fitness and excluded other *rpfR* lineages (18), and we explore other explanations below. In contrast, *ΔrpfF* and WT fail to invade each other when rare (Fig. S4A), which is consistent with complementation by BDSF produced by the WT competitor that activates the PDE in the deletion mutant. In summary, different *rpfR* genotypes that avoid BDSF-mediated dispersion in various ways readily displace the WT ancestor in our biofilm model and can coexist for hundreds of generations by NFDS.

### Biofilm ecology: (ii) co-aggregation and synergistic interactions

Sustained coexistence of different genotypes in biofilms could be enabled by forming aggregates of different composition and form. We tested this potential mechanism of niche differentiation using fluorescently labeled genotypes to measure their co-localization and total volume by confocal microscopy (Table S5). When cultured separately, both A106P and Y355D formed large, thick aggregates that were well dispersed (Fig. 4A), whereas *ΔrpfRF*+93 produced thinner, more uniform biofilms arranged in small clusters. This result shows that the loss of the RpfRF complex and/or BDSF production alters the form of biofilm development, whereas the point mutants appear to produce larger clusters than those produced by WT (Fig. 4AB, Table S5 and Fig. S4A). Different genotype combinations produced aggregates of varying size and biofilm thickness (Fig. 4B). The difference in biofilm development by *ΔrpfRF*+93 was even more apparent when this mutant was co-cultured with either Y355D or A106P, resulting in thinner, more uniform structures and indicating a dominant effect of *ΔrpfRF*+93 on biofilm development (Fig. 4AB and Table S5). Incidentally, we observed that *ΔrpfF* and *ΔrpfR* formed small clusters when mixed with other mutants, but formed larger aggregates when grown together, which is consistent with cross-complementation (Fig. S4A).

**Fig. 4.**
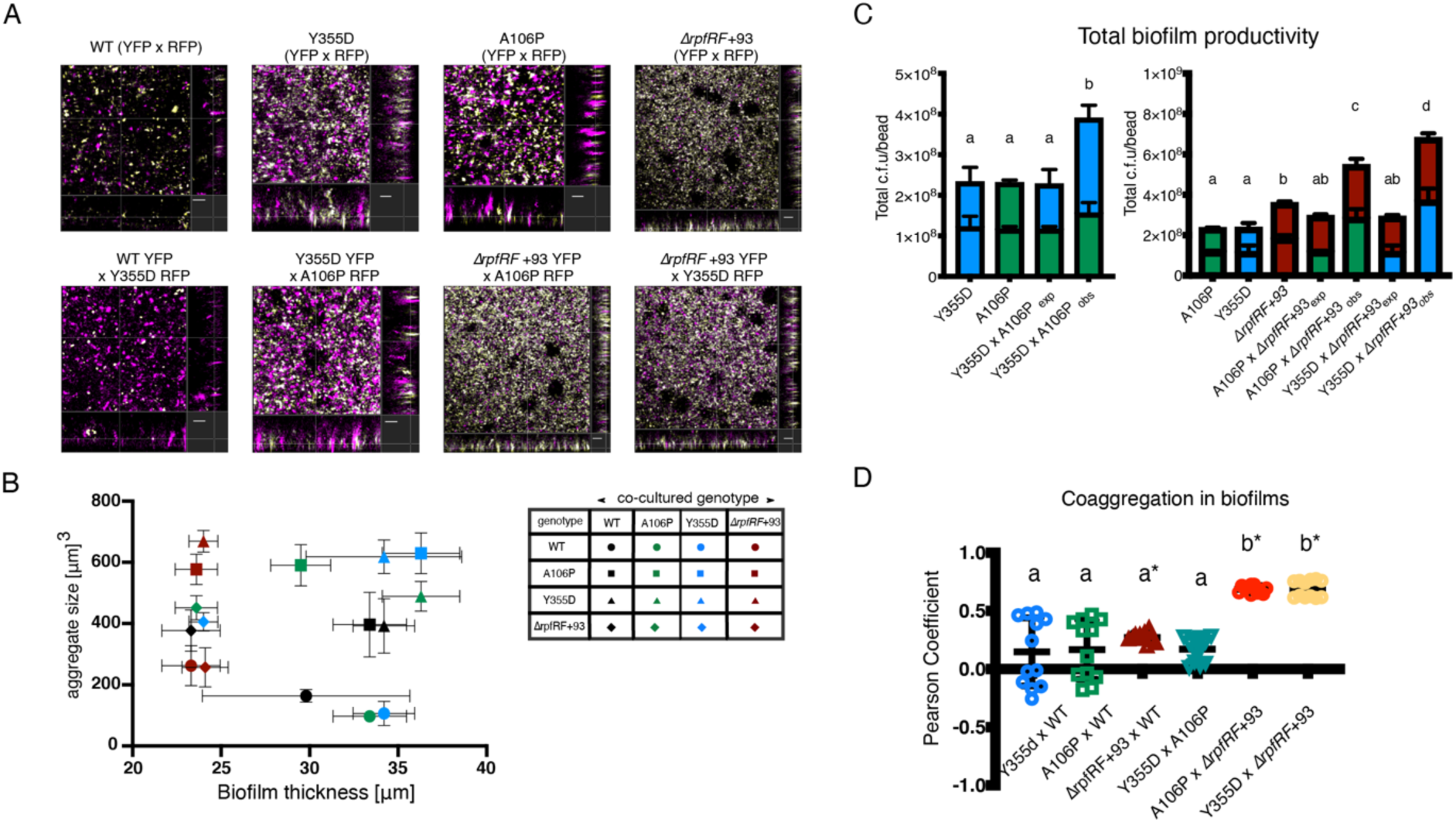
Cocultures of evolved mutants exhibit complementary interactions. (A) Confocal images show structural differences between single strain and coculture biofilms. Large, dispersed aggregates are seen for Y355D, A106P, and Y355D x A106P; monocultures of *ΔrpfRF*+93 and its cocultures (A106P x *ΔrpfRF*+93, and Y355D x *ΔrpfRF*+93) produce small clusters and uniform thickness. RFP-labeled cells are false-colored in magenta and YFP-labelled cells in yellow. White spots indicate coaggregation of differently labeled strains (scale = 10μm). (B) Correlation between average aggregate size of attached aggregate and biofilm thickness. (C) Total biofilm productivity as CFU of individual strains in coculture. A106P, green, Y355D, blue, *ΔrpfRF*+93, red. Expected (exp) values are projected from the individual competitions while the observed (obs) values are experimentally determined. Letters denote significant pairwise statistical groupings. (D) Coaggregation in biofilms, where positive coefficients indicate the extent of overlap between two channels (values significantly different than 0 are denoted with *).

Interactions between genotypes can range from antagonistic, which would reduce net productivity of both types, to synergistic, which would increase productivity of both. We measured productivity as attached CFU/ml and microscopic biovolume for all genotype combinations. In most cases, co-cultures of evolved *rpfR* mutants grown on polystyrene beads were significantly more productive than mutants grown alone (Fig. 4C and Table S5). This indicates that different *rpfR* genotypes facilitate attachment and growth of one other, which supports conclusions from prior studies of long-term evolved biofilm populations (18). Notably, the biofilm productivity of co-cultures of Y355D and A106P is higher than that of the individual genotypes but is lower than either co-cultured with *ΔrpfRF*+93, with increased coaggregation with both point mutants (Pearson coefficient >0.5) (Fig. 4CD, Table S5). These results demonstrate that mixtures of *rpfR* mutants that vary in c-di-GMP levels and BDSF signaling capacity are more productive than when grown alone and produce more uniform biofilm structures together. We speculate that the increased evenness of the mixed biofilm architecture may be an adaptation to maintain attachment to the polystyrene beads, which collide frequently in the test tubes.

### Biofilm ecology: (iii) Polysaccharide composition

*B. cenocepacia* encodes the capacity to produce various polysaccharides. The best known of these is cepacian (composed of rhamnose, mannose, glucose, galactose and glucuronic acid) (28) but others include Bep (*Burkholderia* extracellular polysaccharide) and galactan-KDO (29–31). We hypothesized that the different binding and aggregation properties of *rpfR* mutants related to production of the components in these exopolysaccharides of varied composition. We used fluorescein-tagged lectins that bind different sugars to visualize and quantify differences in the EPS composition of evolved mutants (32). All genotypes including WT produced a matrix composed of mannose, and this sugar was particularly elevated in the *ΔrpfRF*+93 genotype. However, fucose was only detected in *rpfR* mutants, and not *ΔrpfF* (Fig. 5B). Galactose, N-acetyl glucosamine and N-acetyl galactosamine were not detected in the EPS produced by any genotype (data not shown). We then used calcofluor white to stain cellulose and found that Y355D produces much more cellulose than any other mutant (Fig. 5C and Fig. S5). Thus, the varied biofilm phenotypes of *rpfR* mutants may result from secreting different polymers that could serve as shared products that benefit collective attachment.

**Fig. 5:**
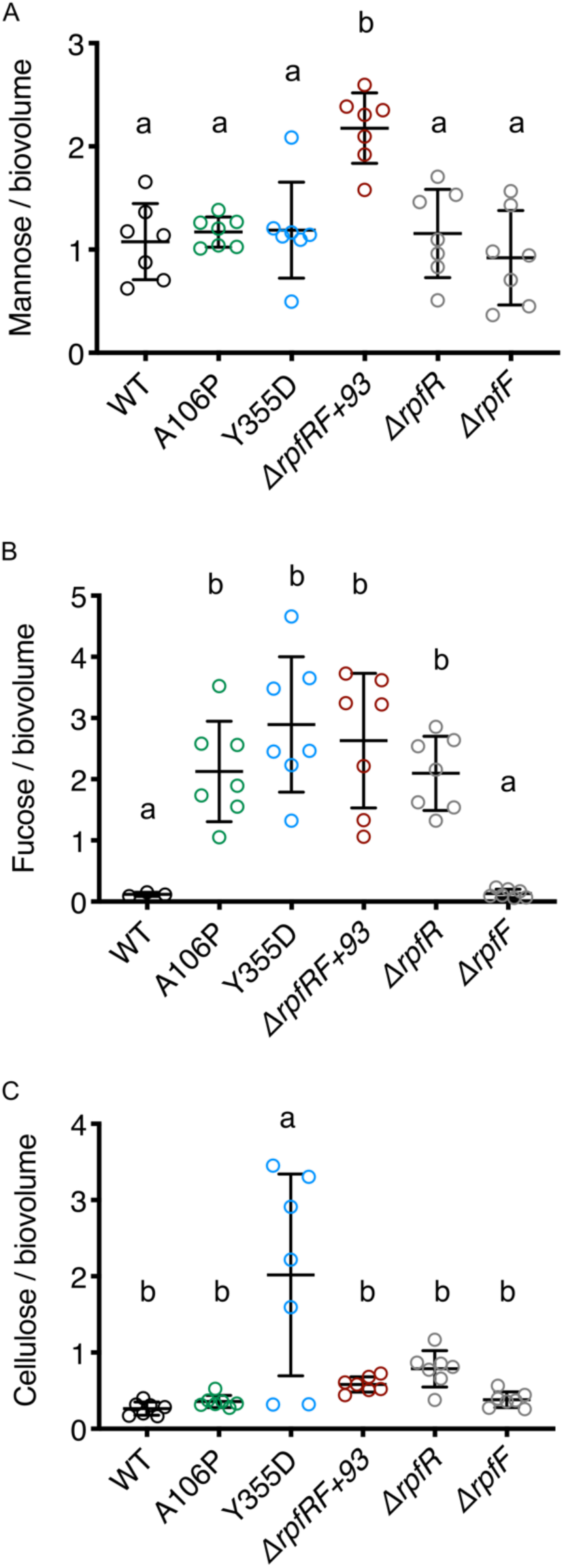
Varied exopolysaccharide (EPS) composition of evolved *rpfR* mutants. Biovolumes were calculated from the fluorescent intensities of bound fluorescently tagged lectins, relative to total bacterial volume, for (A) mannose and (B) fucose or calcofluor for (C) cellulose, and labeled cells using IMARIS 9.0. Different letters indicate significant differences between the mutants.

### Transcriptomic differences among *rpfR* mutants

Mutations in *rpfR* are clearly pleiotropic so to examine the extent of their altered regulation we conducted RNA-seq of six genotypes (A106P, Y355D, *ΔrpfR, ΔrpfF, ΔrpfRF*, and WT) grown under selective biofilm conditions. Hundreds of genes distinguished mutant expression from WT (at q values < 0.05), with Y355D recording the greatest number (∼ 930 genes at Fold change < 1.5) and dozens of genes separated mutants from one another (Fig. S7). As expected from the elevated c-di-GMP levels of mutants, motility and chemotaxis processes were downregulated (except not in *ΔrpfR*, which also produced near-WT levels of c-di-GMP), and in the mutant with the highest c-di-GMP levels, Y355D, other PDE’s (e.g. Bcen2424_5027) were upregulated (Fig. 6). One gene cluster encoding the synthesis of Bep exhibited the greatest increase in expression across all mutants, which provides strong evidence that this polymer is responsible for increased biofilm production. Further, the *berA* gene (Bcen2424_4216), which binds c-di-GMP and activates Bep production and cellulose synthesis, was upregulated in all mutants but *ΔrpfRF* (29, 33). Notably, the genes within the Bep cluster show variable expression levels among mutants, with the most upregulated being the Bcen2424_4206 gene (a *manC* homolog) that encodes mannose-1-phosphate guanylyltransferase. This enzyme plays dual roles, acting as a transferase to convert mannose-1-phosphate to GDP-mannose, a precursor for other sugar nucleotides such as GDP-fucose and GDP-rhamnose, and as an isomerase on mannose-6-phosphate to produce fructose-6-phosphate for gluconeogenesis (34). We hypothesize that increased expression of this gene may activate fucose synthesis (Figure 5) via the intermediate GDP mannose. Interestingly, both *berA* and *manC* show the highest fold-expression changes in Y355D, which could explain the high fucose and cellulose in the EPS of this mutant. Another upregulated gene in Y355D is predicted to encode Flp/Fap pilin (Bcen2424_5868, Fig. S7), which is known to initiate surface attachment in many bacteria (35). These differences strongly suggest a genetic basis of functional differentiation among *rpfR* mutants via c-di-GMP-responsive transcription.

**Fig. 6.**
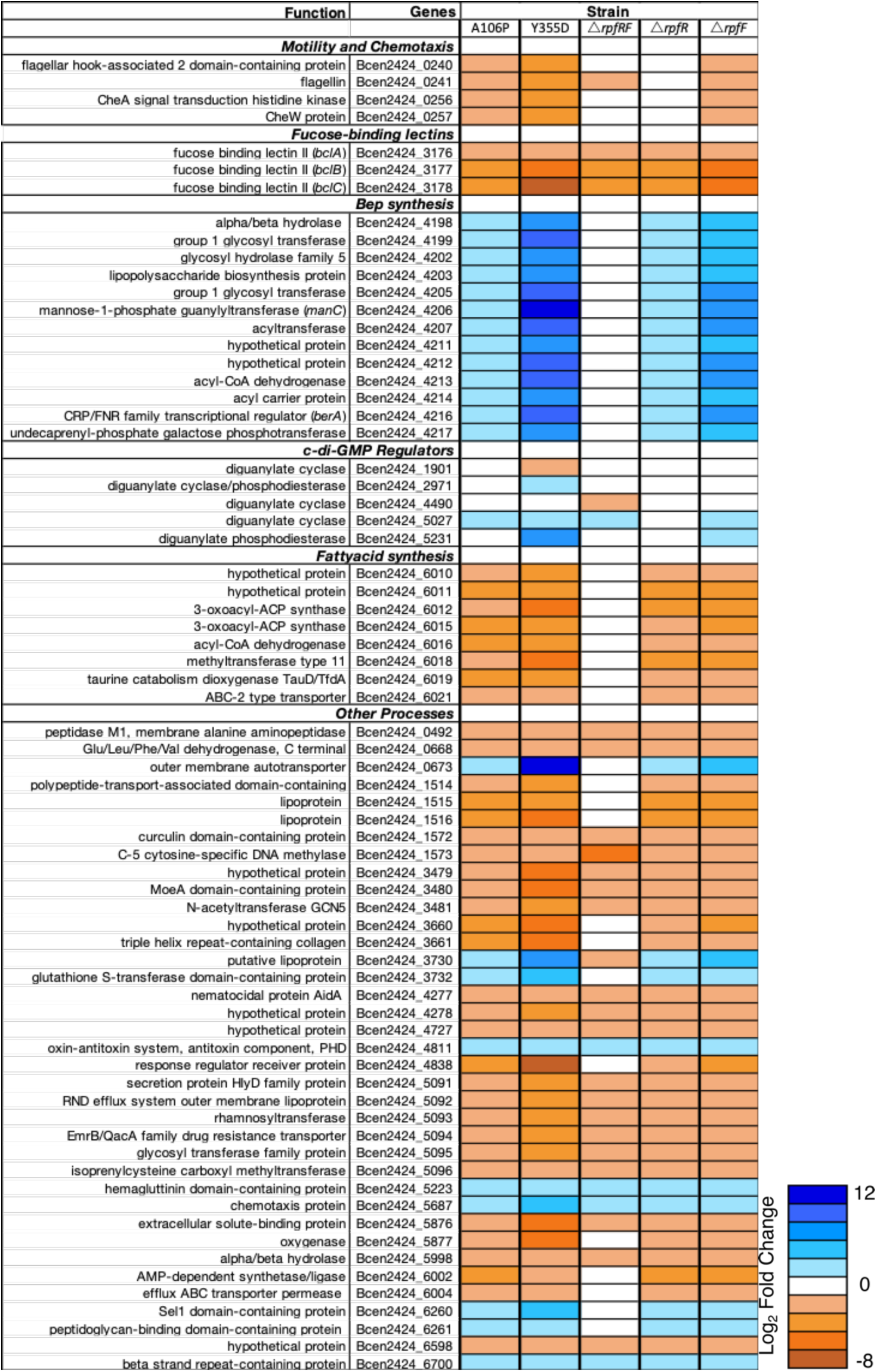
Global changes in expression in evolved and engineered *rpfR* and *rpfF* mutants grown in biofilms. Genes that differentiated 4 or 5 mutants from WT are shown and categorized by function (q value <0.05). Upregulated and downregulated processes are plotted in shades of blue and orange, respectively. Results are from three biological replicates and were analyzed as described in Methods.

The gene cluster most consistently downregulated among *rpfR/F* mutants encodes three fucose-binding lectins (36). These lectins reportedly have high affinity for galactose and fucose and bind carbohydrates in mucus or glycoconjugates at epithelial cell surfaces, which enables them to adhere specifically to host surfaces as single cells (37), but this form of attachment is unavailable in our laboratory system. This result also indicates that *rpfR/F* balances solitary lectin-based attachment against aggregate formation via polysaccharide synthesis. Another cluster that was downregulated among *rpfR/F* mutants putatively encodes fatty acid biosynthesis (Fig. 6). Overall, selection appears to have favored these *rpfR* mutants because of their global regulatory effects that produce a variety of phenotypes related to attachment and biofilm production, as well as dispersal and reattachment. While many of them can be explained generally as classic outcomes of high c-di-GMP, mutants are also differentiated in their patterns of expression. Importantly, demonstrating the power of evolution as a forward genetic screen, the *rpfR* deletion produced the least number of expression changes (Fig. 6), this deletion did not evolve in our experiments, and this mutant was less beneficial than the evolved SNPs that changed but did not eliminate RpfR function.

## Discussion

Many microbes living at surface-liquid interfaces undergo a cycle of attachment, biofilm assembly, dispersal, and reattachment, and thus experience chronic heterogeneity. At the start of these evolution experiments we anticipated diverse genotypes producing adaptations to subsets of these conditions (20, 38). However, much to our surprise, mutations in one gene were selected far more often than any other (13, 18–21). The evolution experiments summarized here collectively span >20,000 generations, yet mutations in only one of the 25 genes in the *B. cenocepacia* HI2424 genome with the DGC or PDE domains that synthesize or degrade c-di-GMP reached high frequency. This focused selection on *rpfR* and the remarkable parallelism at few residues (Fig. 1B) demonstrates that it is the central regulator that governs the switch to biofilm growth. More surprising, because *rpfR* mutations were often the first to reach high frequency in evolved populations, we can infer that only one gene of the predicted 6812 in the *B. cenocepacia* genome encodes the latent potential for the best adaptations in our laboratory biofilm system. This parallelism is at least partly a product of our strain choice and specific experimental conditions, but nonetheless, we expect that *rpfR* plays a similar central role in many other species, where this gene is very well conserved (>60% identical and >80% similar) across dozens of beta- and gamma-Proteobacteria genera and is often syntenic with *rpfF* (Supplemental Data) (12, 39). Our evolution experiments have identified a regulator at the core of c-di-GMP signaling and life history decision-making for numerous bacterial species including many of medical and agricultural significance.

### Diverse effects of evolved mutations extend the model of RpfR / RpfF regulation

Findings of molecular parallelism in evolution experiments are becoming more common and can indicate the functional importance of certain residues. Here, we observed residue-level parallelism at Y355 and R377 in the DGC domain and S570 and F589 in the PDE domain, as well as disproportionate numbers of mutations in the linker regions that connect binding and catalytic domains of RpfR (Fig. 1B). Note that Y355 is 99% identical and R377 is 97% identical across homologs, providing evidence of their functional importance (Table S1). Together, these results demonstrate that selection increased biofilm-related fitness by altering the regulation of but not eliminating RpfR functions. We anticipate these residues are significant for understanding how RpfR, as the first reported c-di-GMP-regulator that is directly activated by a diffusible autoinducer, coordinates diverse responses (11).

Contrary to an earlier report (40), we found that deleting *rpfR* did not cause a growth defect but rather increased fitness in our biofilm model and decreased motility (Fig. 2 and 3). Likewise, deletion of the homolog *pdeR* (previously, *yciR*) in *E. coli* also reduces motility (24). We conclude that RpfR is mainly a PDE with constrained DGC activity, and we speculate that Y355 and R377 play an important role in a conformational change that either activates or inhibits RpfR DGC activity. Likewise, S570 and F589 are 100% identical across *rpfR* homologs and comprise a conserved “loop 6” domain that enables dimerization of the EAL domain and binding of c-di-GMP and the magnesium ion cofactor (41). This study of loop 6 showed that S570 in particular is essential for c-di-GMP binding for hydrolysis, so a point mutation at this site almost certainly enables maintenance of high c-di-GMP levels.

We also found that RpfR interacts directly with RpfF, the enzyme that synthesizes BDSF, and that the RpfR-RpfF interaction inhibits BDSF synthesis (12), We hypothesized that BDSF, RpfR, and RpfF could form a feedback inhibition apparatus whereby BDSF binding to RpfR limits RpfF activity, i.e., BDSF production. Further, we predict that the RpfR-RpfF interaction is critical in the long term for these bacteria but dispensable in these short-term experiments. RpfF synthesizes BDSF by dehydrating 3-hydroxydodecanoyl-acyl carrier protein (ACP) to form *cis*-2-dodecenoyl-ACP, and hydrolyzing the thioester bond linking the acyl-chain to ACP, releasing free BDSF (42). However, RpfF is promiscuous and can target other acyl-ACP substrates, hampering membrane lipid synthesis for example. Some bacteria like *Xanthomonas spp*. also produce antagonist proteins RpfB and RpfC to control RpfF activity (40), but *Burkholderia* lacks these proteins. Thus, the RpfR-FI domain is key to governing RpfF activity. This regulation is in addition to the interaction between BDSF and RpfR, which activates its PDE domain upon binding the PAS domain (11) (Fig.1).

Building upon this model, we predict that the parallel A106P mutation in the linker region between the FI and PAS domains interferes with a conformational change that activates the PDE domain upon BDSF binding (Fig. 7). Mutants in this linker can be considered “signal-blind” and maintain basal DGC activity, which is consistent with the intermediate c-di-GMP and fitness effects of this mutant (Fig. 2 and 3). Another common mutation completely deleted *rpfR* and *rpfF* and 93 other genes, which eliminates both BDSF synthesis and RpfR-mediated regulation of c-di-GMP by its dominant PDE. This should lead to a net increase in biofilm production and biofilm-related fitness, which we observed, but also an inability to either produce or sense BDSF and thus a relative insensitivity to the functions of other genotypes. This predicted signal-blind and -mute function is consistent with the ability of this genotype to persist and ultimately invade the other mutants with the benefit over other mutations in the LTE (13), despite its lower initial fitness observed.

**Fig. 7.**
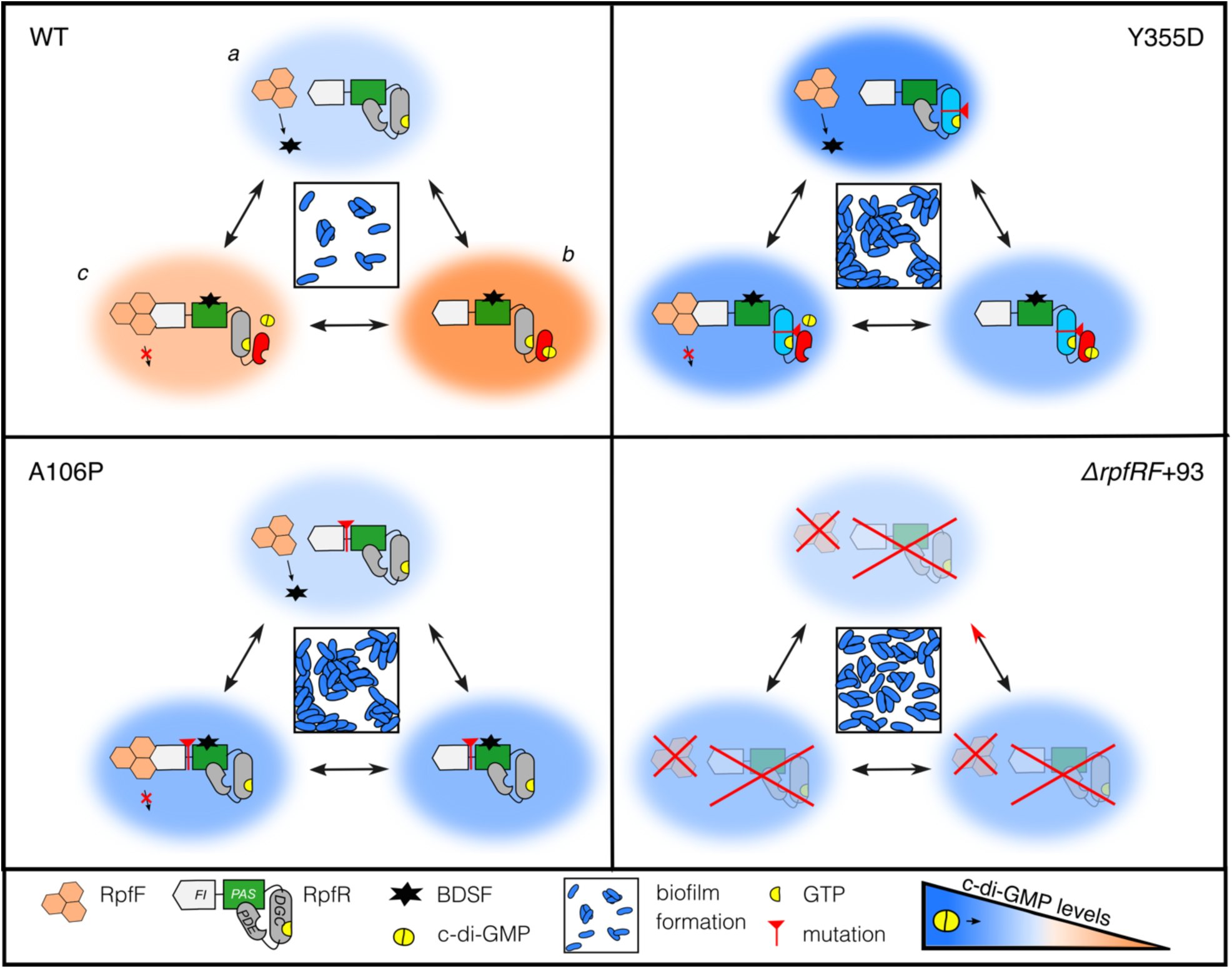
Predicted effects of biofilm-adapted *rpfR* mutants on the RpfR/F signaling complex. Top left: the WT genotype produces sparse biofilms owing to its production (a) and sensing of BDSF, which binds RpfR-PAS and activates the RpfR-EAL phosphodiesterase domain that hydrolyzes c-di-GMP (orange color gradient, b). The RpfR-FI domain limits RpfF activity by binding and inhibiting BDSF synthesis, enabling c-di-GMP levels to recover to low levels (c). Top right: the Y355D genotype hyper-activates the GGDEF domain and increases c-di-GMP, resulting in large biofilm aggregates. Although BDSF binding to RpfR-PAS can activate the phosphodiesterase domain, c-di-GMP levels remain high (blue color gradient). Bottom left: we hypothesize that A106P in the linker region between the FI and PAS domains prevents a conformational change caused by the BDSF-PAS interaction, rendering this genotype blind to BDSF. Hence, c-di-GMP levels increase slightly either by the action of RpfR or other DGCs. This mutant also forms large biofilm aggregates. Bottom right: In the absence of both RpfF and RpfR, no BDSF is produced and c-di-GMP levels produced by other enzymes accumulate, producing a biofilm composed of small aggregates. This phenotype is dominant to the other genotypes, and mixtures take on this more uniform biofilm phenotype.

Integrating these findings allows us to expand our mechanistic understanding of how this RpfF/R regulatory node governing c-di-GMP signaling and BDSF quorum sensing enables “decisions” within the biofilm life cycle (Fig. 7). Conditions that select for increased biofilm would favor deactivation of PDE activity either directly, by mutating S570 / F589, or indirectly, by limiting BDSF binding to activate the PDE (A106P) or by eliminating BDSF synthesis by RpfF (Δ*rpfF*). Alternatively, mutants like Y355D that activate the DGC would be selected (Fig. 7). We tested these predictions by making targeted mutations of the functional domains of this system. First, in a prior study we engineered point mutations in the PAS domain at sites predicted to bind BDSF, and these produced elevated c-di-GMP and fitness because the PDE domain was not activated (12). Here, we also deleted the FI domain thought to control RpfF activity and, as expected, this mutant had low c-di-GMP and was deleterious under biofilm conditions (Fig. 2CD and 3A). On the other hand, deleting *rpfF* greatly increased c-di-GMP, but curiously this single-gene deletion was not beneficial in competition with WT, perhaps because the WT complemented the BDSF defect of Δ*rpfF* in cocultures, and consequently was never selected in our experiments (Fig. 3). This implies that the RpfR-F complex, perhaps also with other partners (9), has been preserved by selection as a functional unit and that disrupting only one component is disfavored. The Hengge laboratory has advanced the model that the RpfR ortholog in *E. coli*, PdeR, functions as a “trigger enzyme” at the hub of c-di-GMP signaling to control curli synthesis and other biofilm-related traits (43). Although *E. coli* does not encode RpfF, it is possible that the FI domain of PdeR and its orthologs in diverse species bind other proteins contributing to the trigger.

### Ecological diversification and complementary lifestyles are pre-wired within RpfF/R

In retrospect, perhaps we should not have been surprised that mutants of a multi-domain protein with both sensory and catalytic activities would have varied functions. What is remarkable is that different mutants can evolve and coexist in the same populations because of their distinct ecological consequences. For instance, the small aggregate phenotype of *ΔrpfRF*+93 allows growth between the large aggregates of its co-cultured partner, increases overall biofilm productivity, and maintains genetic diversity despite the dominant Y355D genotype (Fig. 4). These mixed biofilms consisting of genotypes producing large aggregates and small clusters appear to decrease competition and increase the carrying capacity of the environment, which is consistent with the character displacement process we described previously in a long-term evolution experiment (18, 44). Overall, higher c-di-GMP levels correlated with more EPS production and larger aggregates, but new biofilm phenotypes emerged when evolved mutants were co-cultured (Fig. 4), including reciprocal frequency-dependence (Fig 3). These positive interactions explain how three different *rpfR* mutants were maintained for hundreds of generations in the first long-term experiment (13).

The phenotypes encoded by this system contribute to the important differences in affinity for different surfaces and potential for pathogenesis. While the WT strain produces the potent BclACB lectins that bind fucose and mannose residues on host cells (Fig. 6) (45), the evolved mutants downregulate these lectins to upregulate EPS production, whose composition varied among mutants (Fig. 5). EPS synthesis is associated with upregulation of the *manC* gene, which is reported as a virulence factor in cystic fibrosis infections caused by the *Burkholderia cepacia* complex (46). Lectin synthesis is under control of quorum sensing molecules including BDSF (47), and also by the protein GtrR that binds to the *bclABC* promoter and induces their expression. RpfR enhances this expression by forming a complex with GtrR, but not when it binds c-di-GMP (14). It follows that the high c-di-GMP production of evolved *rpfR* mutants downregulated lectin production. However, these mutations encumber a tradeoff that limits other dimensions of the niche, like reduced motility and suppressed lectin-based attachment (Fig 5 and Fig. S1B and S5), and likely would not persist over the longer term in nature.

The *Burkholderia cepacia* complex is best known for causing opportunistic infections in the cystic fibrosis airway, where populations encounter a more restrictive subset of their original niche that selects for traits like aggregation regulated by *rpfRF* (19). Tracking evolving populations of species with *rpfRF*, either *in vitro* or *in vivo*, will provide valuable tests of the model presented here and determine whether *rpfRF* is a c-di-GMP signaling node in other species that could eventually be exploited for antimicrobial strategies or microbiome engineering.

### Experimental Procedures (to appear in SI)

#### Bacterial growth media and conditions

Strains and plasmids used in the study are listed in Table S2. The frozen stocks were revived in Tryptone Soy Broth (TSB) and preconditioned in 3% galactose M9 minimum medium, hereafter GMM (17) for all the experiments unless specified otherwise. For GFP and RFP-labeled strains, 100 µg/ml concentration of chloramphenicol antibiotic was supplemented in the growth media. TSB agar was used for enumerating colony forming units (CFU) and culturing purposes. X-gal was added to the agar plates to differentiate between lac+ and lac-strains.

#### Genetic engineering

Isogenic mutants were created using methods described by Fazli et al (48). Briefly, single and double gene deletions were created by amplifying approximately 1000 bp upstream and downstream of the target gene and then joined using single overlap extension PCR using primers GW-attB1 and GW-attB2 (Table S2). The resulting approximately 2000 bp fragment was then inserted into a pDONPREX18Tp-SceI-PheS plasmid using Gateway cloning. For single-nucleotide mutations, the target gene was amplified, cloned into the above plasmid, and site-directed mutagenesis was used to create the intended point mutations. The resulting vectors were transformed into *E. coli* DH5α, and then into *B. cenocepacia* by conjugation using tri-parental mating previously described (49). Mutants were then selected by sensitivity to 100 µg/ml trimethoprim and sequenced using whole genome sequencing on an Illumina NextSeq 500 to a minimum coverage of 30x (50). We used the variant calling program Breseq v. 0.31to confirm isogenic mutants to be used in the study (51).

To enable competitors to be distinguished in mixed culture, a *lacZ* marker was added using plasmid pCElacZ, as previously described using four parental conjugation (52). For confocal microscopy, WT and mutants were electro-transformed (53) with plasmids pSPY, which harbors yellow fluorescent protein genes and pSPR that contains red fluorescent protein from DsRedExpress (17) and were selected on TSB agar plates containing 100 µg/ml chloramphenicol.

#### Fitness assay

The optical densities of the GMM preconditioned cultures were standardized, and competitors (lac+ and lac-) were mixed in defined ratios (1:1 for direct competition, and when testing for frequency-dependent interactions, at 1:9, 3:7, 7:3 and 9:1 ratios) into 5 replicate tubes containing 5 ml of GMM medium + three 7 mm polystyrene beads each. For planktonic fitness assays, culture tubes contained no beads. An aliquot of that mixture was serially diluted and plated to get the starting CFU followed by incubation of tubes at 37°C, shaking conditions. At 24 h, one bead was transferred using ethanol-sterilized forceps to a new culture tube containing two different colored beads to determine fitness at 48h while another was sonicated using a probe sonicator at a continuous pulse for 10 secs at 30% amplitude. In the case of planktonic growth, a planktonic fraction was sampled for the CFU counts and for the inoculation in the new tube for 48 h measurement. The 24 and 48 h samples thus collected were serially diluted and plated on TSB X-gal plates. Selection rate was calculated as the difference in the Malthusian parameters of the two competitors using the equation, s= ln[A(t)/A(0)] – ln[B(t)/B(0)], where A and B are the two competitors quantified at time 0 and t. The values were graphically represented and statistics were performed on Graphpad Prism 8. The Two-stage linear step-up procedure of Benjamini, Krieger, and Yekutieli, q value< 0.05 was used to perform pairwise comparisons post-hoc test.

#### Quantification of cellular c-di-GMP

1.25 mL of preconditioned cultures were added to 125 mL of GMM in flasks containing 100 7 mm polystyrene beads each and incubated at 100 rpm at 37 °C for 12 h. While harvesting, flasks were incubated on ice for 10 min. For planktonic phase, 25 ml of the culture was transferred to 50 ml centrifuge tube. For the biofilm phase, the planktonic culture was discarded and the beads were washed with 60 mL of cold PBS. These were then divided into four 50 mL centrifuge tubes containing 20 mL of cold PBS each. Each tube was vortexed for 30 s to remove the attached cells and the PBS from all 4 sets was combined. The samples were serially diluted and plated to enumerate CFU/flask and then centrifuged at max speed for 15 min at 25 °C. Pellets were resuspended in 500 µL of ice-cold extraction buffer (methanol:acetonitrile:dH_2_O 40:40:20 + 0.1 N formic acid). The suspensions were transferred to 1.5 mL microfuge tubes and incubated at −20 °C for 1 h, followed by 95 °C for 10 min. The tubes were then centrifuged to pellet the cell debris. 400 µL of the liquid phase was transferred to another microfuge tube and 16 µL of neutralization buffer (15% ammonium bicarbonate) was added. The tubes were stored at −80 °C. Quantification of cdG using mass spectroscopy was then carried out as previously described (54).

#### Biofilm assay

GMM preconditioned cultures were inoculated (1:100) in 96-well microtiter plate containing 200 µl of GMM per well (the peripheral wells were not inoculated as were used as blank readings). The plate was incubated at 37°C under static conditions. The following day, medium was discarded, wells were washed using PBS and later stained with 0.1% crystal violet dye with subsequent 15 min incubation, as described (55). Ethanol solution (95% EtOH, 4.95% dH 2 O, 0.05% Triton X-100) was added to the wells to de-stain and the solution was transferred to a new plate. Absorbance was measured at 590 nm. The values were then normalized using blank readings and the resultant values were used to plot the graph.

#### Colony morphology

GMM preconditioned cultures were spotted (4 *µ*l) on Congo red Tryptone 0.7% agar plates or serially diluted to plate on TSB 1.5% agar. The plates were incubated at 37°C under static conditions. Next day the plates were placed on the bench to allow structures to develop. The spot-colonies were imaged using Nikon D3300 with 18-55mm lens while the isolated colonies were captured on a brightfield microscope fitted to Canon DS126491 with a 2X microscope adapter lens. Images were scaled using ImageJ.

#### Confocal microscopy

Fluorescently labeled cultures were preconditioned in GMM + 100 µg/ml chloramphenicol to maintain plasmid carriage. Equal volumes of these were added to a 1.5 ml microfuge tube containing 800 µl GMM. The mix was vortexed and 200 µl was transferred to an optically clear bottom 96-well microtiter plate in triplicates. The plate was incubated at 37°C at 100 rpm shaking conditions. The z-stack images of the biofilms formed at the bottom of the wells were taken using Olympus FLUOVIEW FV3000 confocal laser scanning microscope with 20 X objective lens [excitation at 488nm (EGFP) for YFP and 560 nm (TRITC) for RFP). For staining polysaccharides in biofilm, RFP labeled monocultures were inoculated in wells. At 24 h, the supernatant was carefully removed and 50 µl of 50 µg/ml fluorescein-tagged lectins (Concanavalin A for mannose and Ulex Europeus agglutin for fucose, Vector Laboratories) were added (32). To stain cellulose, 1:10 solution of Calcofluor white was added to the wells. After adding the stains, the plates were incubated at room temperature for 20 mins. The stain solutions were removed and wells gently washed by pipetting out the well contents and replacing with PBS. Biofilm images were captured at an excitation of 560 nm for RFP, 494 nm for fluorescein, and 365 nm for Calcofluor white. The image stacks were analyzed using IMARIS 9 for creating orthogonal view images [Center for Biologic Imaging, University of Pittsburgh]. We used the IMARIS extension for Biofilm analysis by Matthew Gastinger to quantify biofilm parameters such as thickness, total biovolume, and biomass for each channel (http://open.bitplane.com/tabid/235/Default.aspx?id=119). The median of volumes calculated after the surface segmentation in IMARIS was used as average aggregate size value for each channel and the sum of the volumes was equal to the total biovolume calculated. Pearson coefficients were calculated using the Coloc function in IMARIS, as the measure of colocalization between two strains, ranging from 1 to −1 (56–58). Different letters are used to indicate significant differences between the data points (paired t-test calculated using the Two-stage linear step-up procedure of Benjamini, Krieger, and Yekutieli, q value< 0.05).

#### Biofilm productivity

The optical densities of the GMM preconditioned cultures were standardized, and competitors (lac+ and lac-) were mixed in equal ratios in 5 replicate tubes containing 5 ml GMM + three 7mm polystyrene beads. The tubes were incubated at 37°C, shaken conditions. At 24 h, the beads were sonicated and cells were plated on TSB X gal plates. The total cfu/ml was calculated from enumerated colonies.

#### Motility

TSB plates containing 0.3% agar were prepared on the previous day. The following day, GMM preconditioned cultures were spotted on the plates with a toothpick. Each plate had a mutant and a WT control. The plates were incubated without turning upside down at 37°C. Colony diameter was measured at 18h and expressed relative to the corresponding WT diameter.

#### RNA-seq

1.25 mL of three independent preconditioned cultures were added to 125 mL of GMM in flasks containing 100 7mm polystyrene beads each and incubated at 100 rpm at 37 °C for 16 hours. For each flask, the suspension was discarded and the beads were washed with 60 mL of cold PBS. These were then divided into four 50 mL centrifuge tubes containing 20 mL of cold PBS each. Each tube was vortexed for 30 s to remove the attached cells and the PBS from all 4 sets was combined. The suspension was centrifuged at max speed for 15 min at 25 °C. 500uL of RNAprotect was added to the final cell pellet followed by RNA extraction with Amresco Phenol Free RNA kits. The RNA was sequenced at Genewiz and the reads were pseudo-aligned to the HI2424 genome using Kallisto version 0.46 (59). Read counts were quantified through Kallisto at 1000 bootstraps per sample. Differential gene expression analysis began by feeding the raw, quantified read counts into edgeR (60). Raw read counts were first normalized to counts per million, and then further normalized using edgeR’s TMM normalization method. Differentially expressed genes were called by invoking edgeR’s Genewiz negative binomial generalized linear model, whereby variance was inferred between treatments’ replicates. Genes that displayed q <.05, |Fold Change| > 1.5 and at least half of the replicates with counts per million > 1 were considered differentially expressed. The genes demonstrating statistically significant fold change with respect to wildtype in at least 4 strains were organized by functional categories using https://www.burkholderia.com. The raw reads are available at NCBI Bioproject PRJNA607303.

#### Statistical analyses

Statistical analyses were conducted in Graphpad Prism 8 for Mac OS X, GraphPad Software, La Jolla California USA, www.graphpad.com, or in R (61). Mutant comparisons were conducted by one-way ANOVA with post hoc testing using the two-stage linear step-up procedure of Benjamini, Krieger, and Yekutieli.

## Acknowledgments

This research was supported by grants NIH R01GM110444 and NASA NAI CAN-7 NNA15BB04A to VSC. We thank Prof. Simon C. Watkins, director of Center for Biologic Imaging (CBI), University of Pittsburgh for IMARIS support and image analysis assistance, members of the Cooper laboratory and Evan Waldron (Rutgers) for helpful discussions and proof reading, and Christopher Deitrick for bioinformatics help and depositing RNAseq files in the NCBI database.

## Supplementary Information

### Supplementary data tables

**Table S1:**
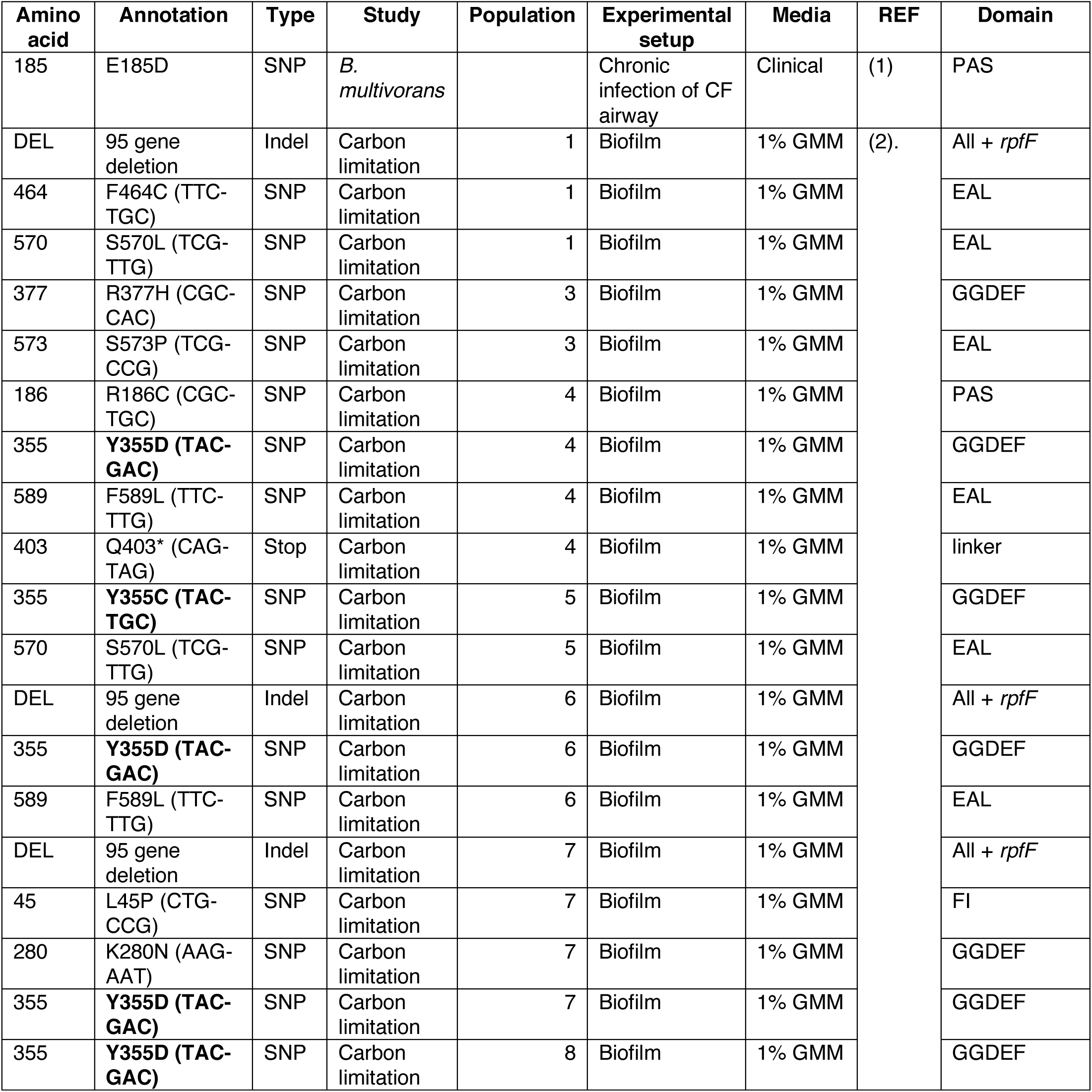
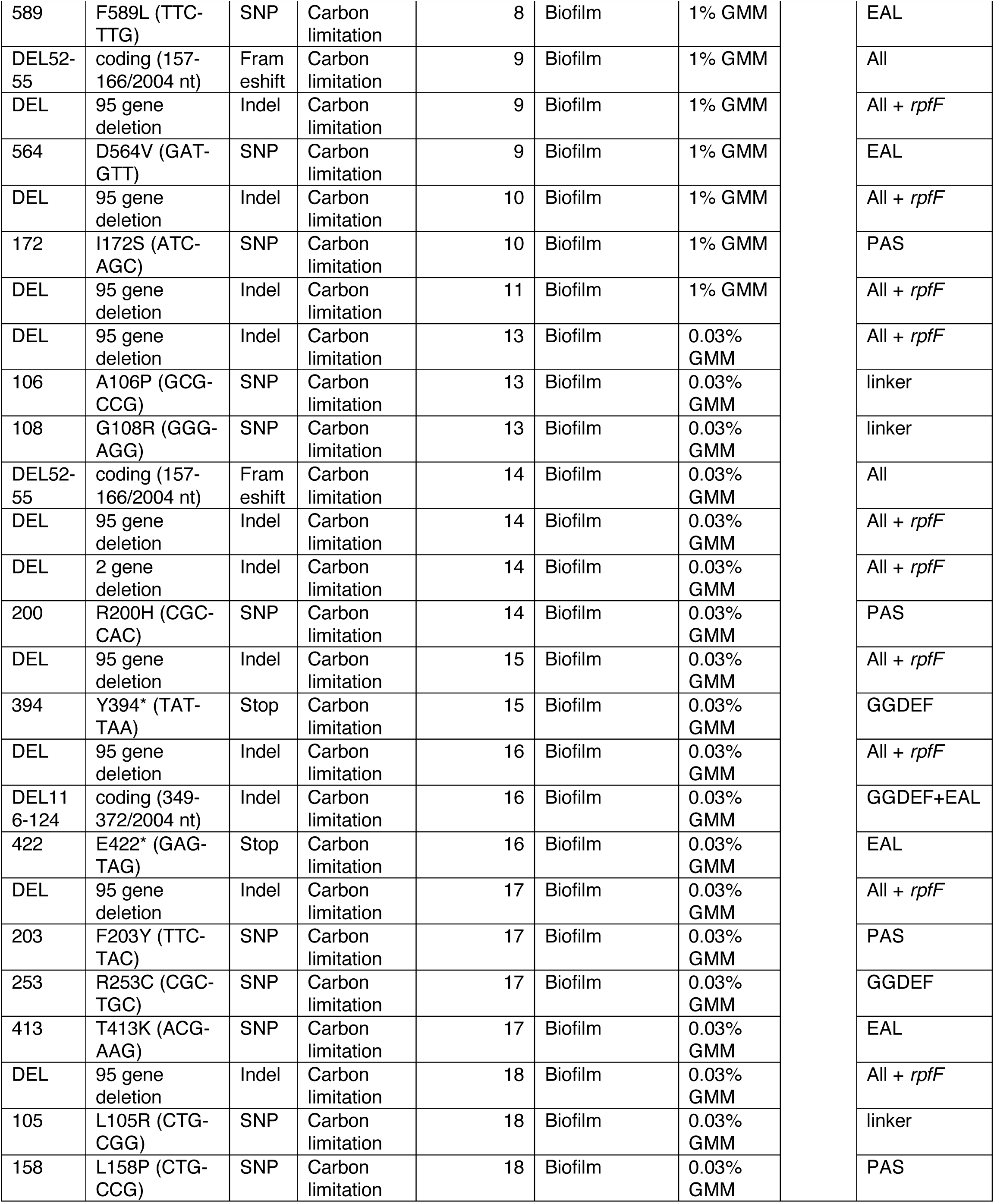
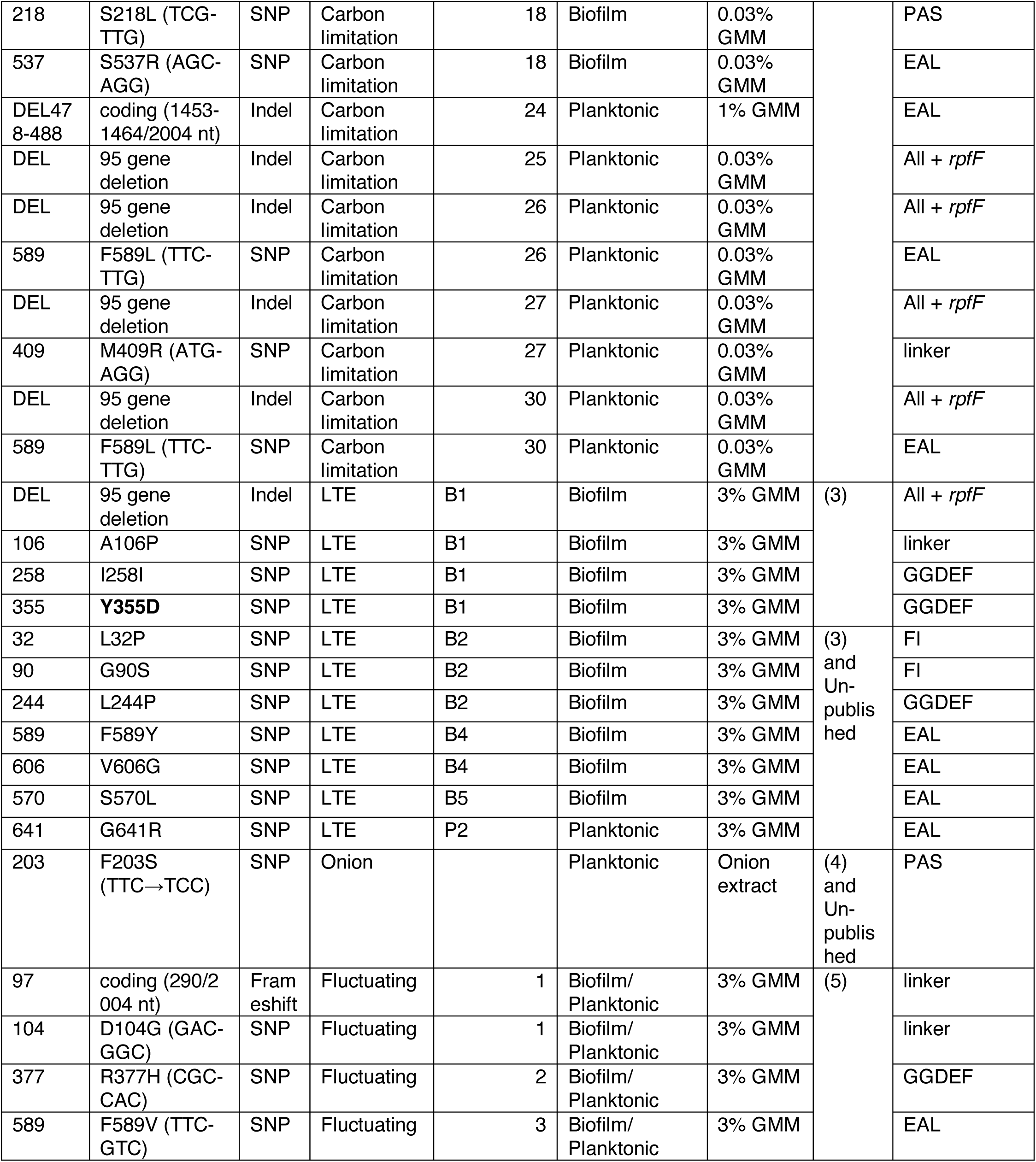
Evolved *rpfR* mutations identified from whole-population genomic sequencing of experimental populations (>5% or greater frequency) or from isolated clones from the experiment or biofilm-associated infection.

**Table S2:** List of bacterial strains used in the study

| Strain | Genotype | Phenotype | Reference |
| --- | --- | --- | --- |
| *B. cenocepacia* |  |  |  |
| WT | HI2424 |  | (6) |
| Y355D | SNP in GGDEF domain of *rpfR*<br>(Bcen2424_3554) | High c-di-GMP,<br>increased biofilm | This study |
| A106P | SNP in the linker region<br>between FI and PAS domains<br>of *rpfR* | Increased biofilm | This study |
| $\Delta rpfRF+93$ | Deletion of *rpfR*, *rpfF* and 93<br>other genes downstream | Increased biofilm | Isolated from population B1<br>(3) |
| $\Delta rpfF$ | Deletion of *rpfF*<br>(Bcen2424_3555) | No BDSF production | This study |
| $\Delta rpfR$ | Deletion of *rpfR* | Increased biofilm | This study |
| $\Delta rpfFR$ | Deletion of *rpfF* and *rpfR* | Increased biofilm | This study |
| GGDAF | SNP in GGDEF domain of *rpfR* | Increased biofilm | This study |
| AAL | SNP in EAL domain of *rpfR* | Increased biofilm | This study |
| FI | Deletion of FI domain in *rpfR* | Increased biofilm | (6) |
| Y355D-GGDAF | SNPs in GGDEF domain of<br>*rpfR* | Decreased c-di-GMP | This study |
| WT lac+ | Tn7::lacZ |  | (4) |
| Y355D lac+ | Tn7::lacZ |  | This study |
| $\Delta rpfRF+93$ lac+ | Tn7::lacZ | | This study |
| $\Delta rpfR$ lac+ | Tn7::lacZ | | This study |
| WT YFP | pSPY |  | This study |
| WT RFP | pSPR |  | This study |
| Y355D YFP | pSPY |  | This study |
| Y355D RFP | pSPR |  | This study |
| A106P YFP | pSPY |  | This study |
| A106P RFP | pSPR |  | This study |
| $\Delta rpfRF+93$ YFP | pSPY | | This study |
| $\Delta rpfRF+93$ RFP | pSPR | | This study |
| $\Delta rpfF$ YFP | pSPY | | This study |
| $\Delta rpfF$ RFP | pSPR | | This study |
| $\Delta rpfR$ YFP | pSPY | | This study |
| $\Delta rpfR$ RFP | pSPR | | This study |
| *E. coli* DH5α | pSPY | | (7) |
| *E. coli* DH5α | pSPR | | |
| *E. coli* DH5α | Tn7::lacZ | | |
| *E. coli* DH5α | pTnHelper | | (8) |

### Supplementary figures

**Fig. S1.**
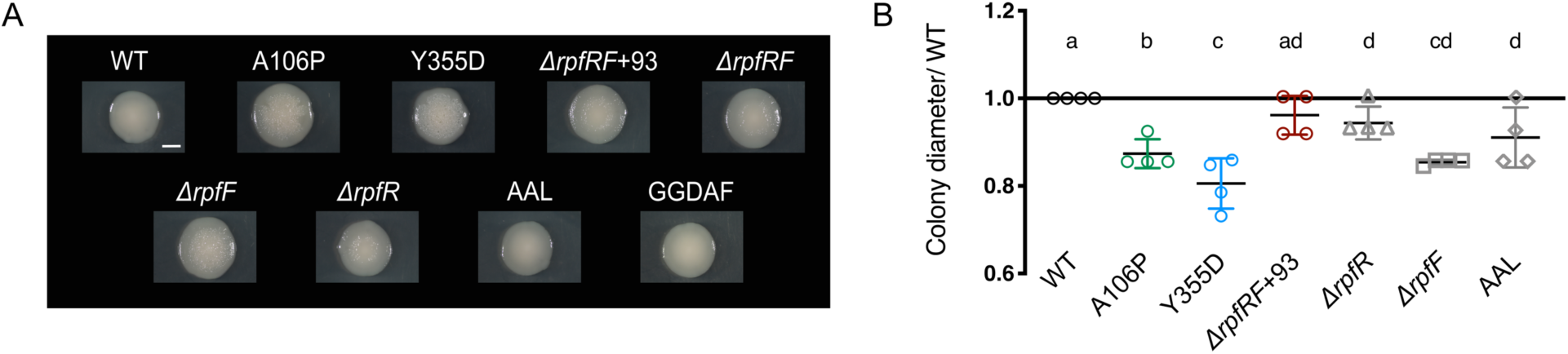
Phenotypic characteristics of *rpfR* mutant colonies. (A) Isolated colonies of *rpfR* mutants (scale bar= 2mm). Spot colonies shown in Fig 1 do not enhance the central studded structures otherwise often seen in isolated colonies on half-strength Tryptic Soy broth. Engineered mutants inactivating the catalytic domains (AAL and GGDEF) have a smooth phenotype. (B) Relative motility is determined by the colony diameter of the mutant divided by that of the wild type after 18 hours of growth in soft agar plates (different letters indicate significant differences between mutants following posthoc Tukey tests).

**Fig. S2.**
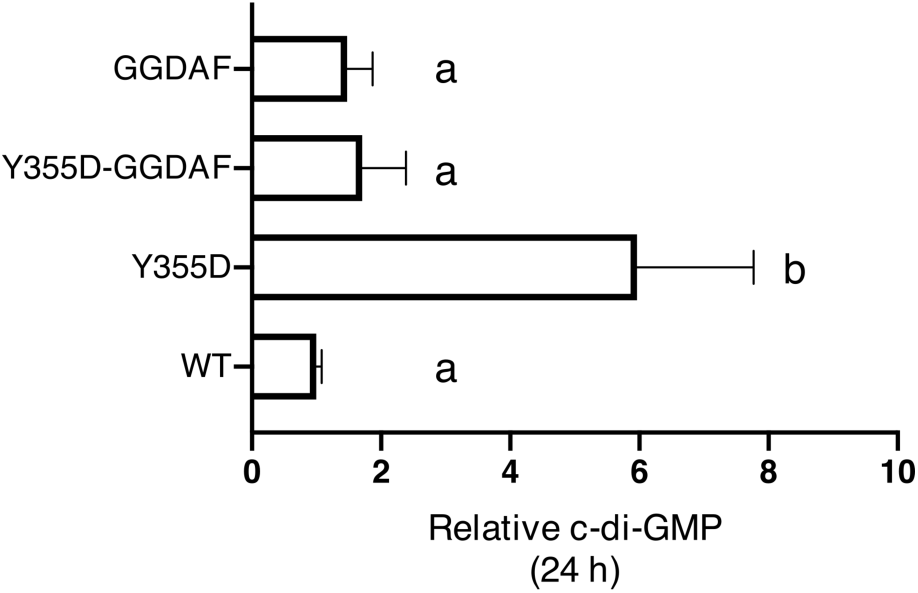
Activation of the GGDEF domain in the Y355D mutant contributes to the enhanced c-di-GMP levels and fitness gain. The c-di-GMP levels in *rpfR* GGDAF mutant inactivating the diguanylate cyclase function are similar to that in WT, indicating that the GGDEF domain in WT is probably inactive. Interestingly, mutation Y355D seems to activate the GGDEF domain suggested by the high levels of c-di-GMP and subsequent lower levels in the Y355D GGDAF mutant.

**Fig. S3.**
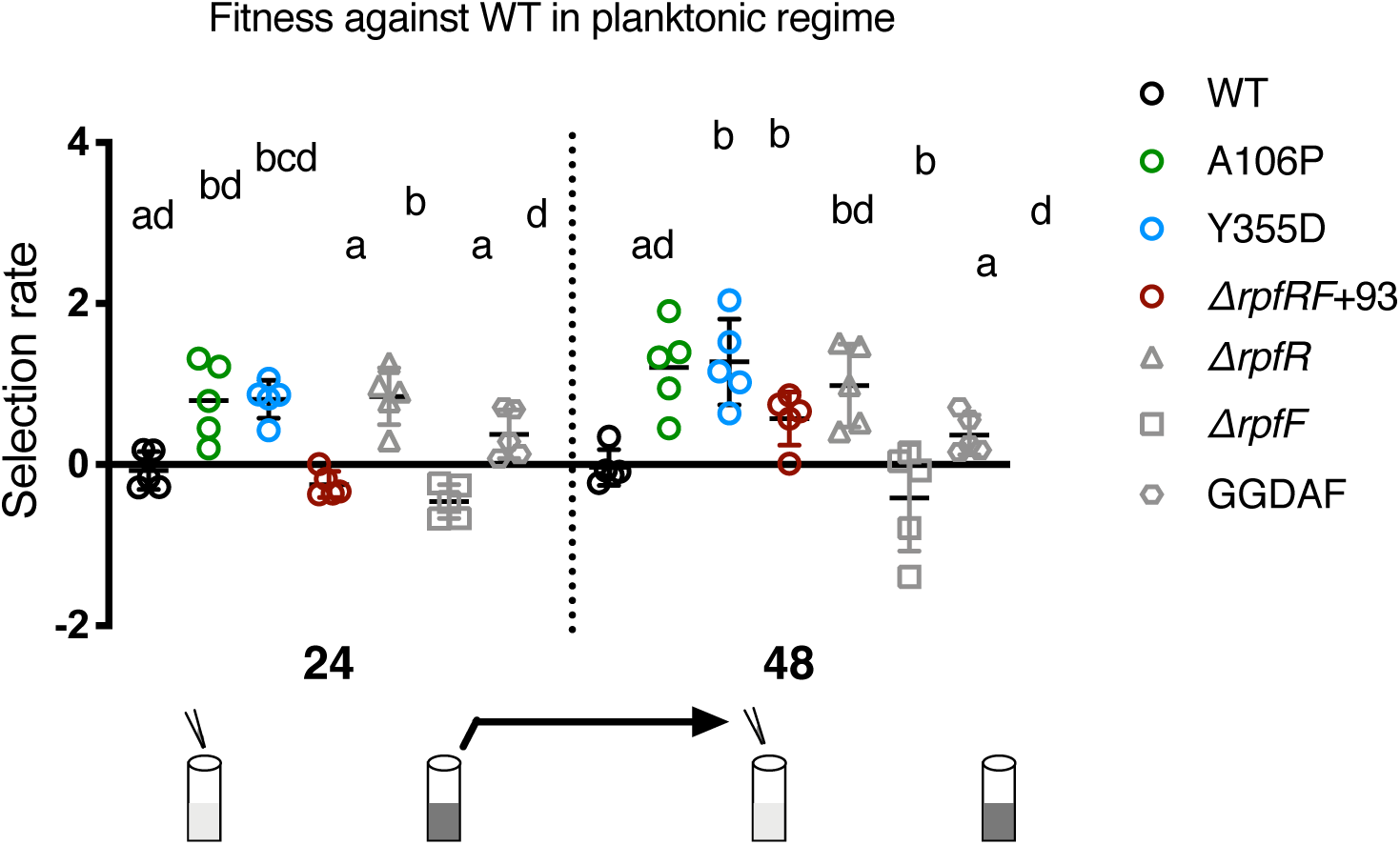
Relative fitness against WT at 24 and 48 hs in planktonic growth conditions. Most mutants exhibit higher fitness in planktonic conditions (except, *ΔrpfRF*+93 and *ΔrpfF*). Different letters are used to indicate significant differences between the mutants by post hoc testing following ANOVA.

**Fig. S4.**
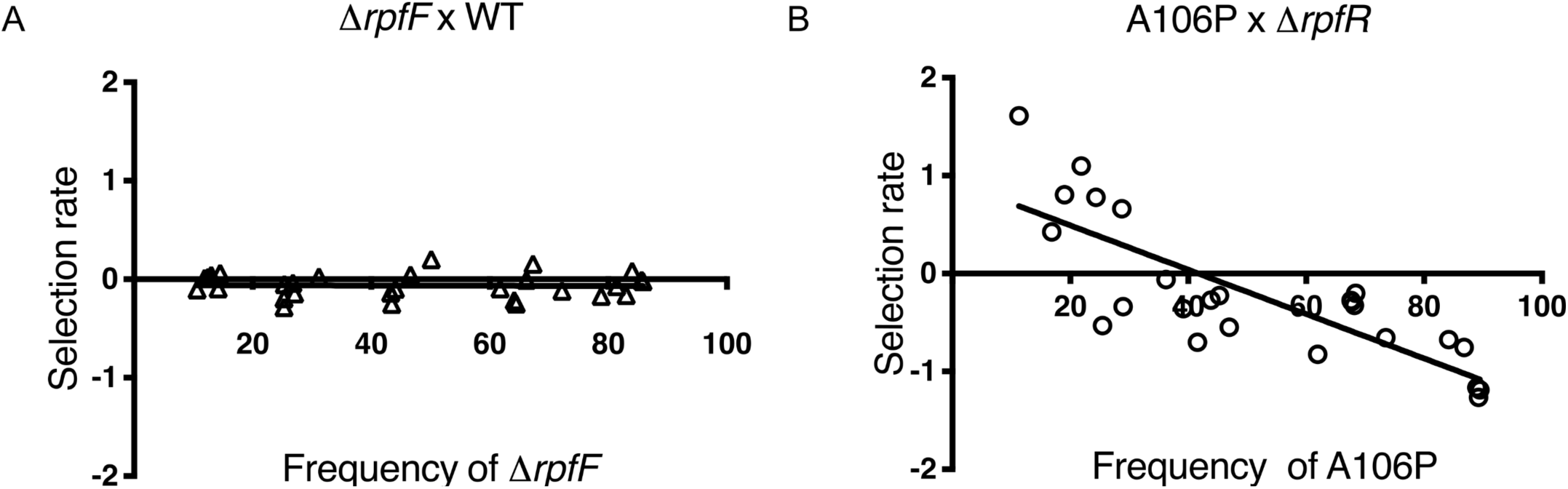
Tests of frequency-dependent interactions. (A) *ΔrpfF* versus WT has no fitness advantage against wild type even when present at different frequencies, despite higher biofilm production when grown alone. (B) A106P versus ΔrpfR, where A106P shows negative frequency-dependent selection against ΔrpfR, consistent with its insensitivity to BDSF production. Regression analyses produced the following functions: y = −0.0001073*x - 0.05973, r^2^= 0.0005545 (A) and y = −0.02265*x + 0.945, r^2^= 0.6421 (B).

**Fig. S5.**
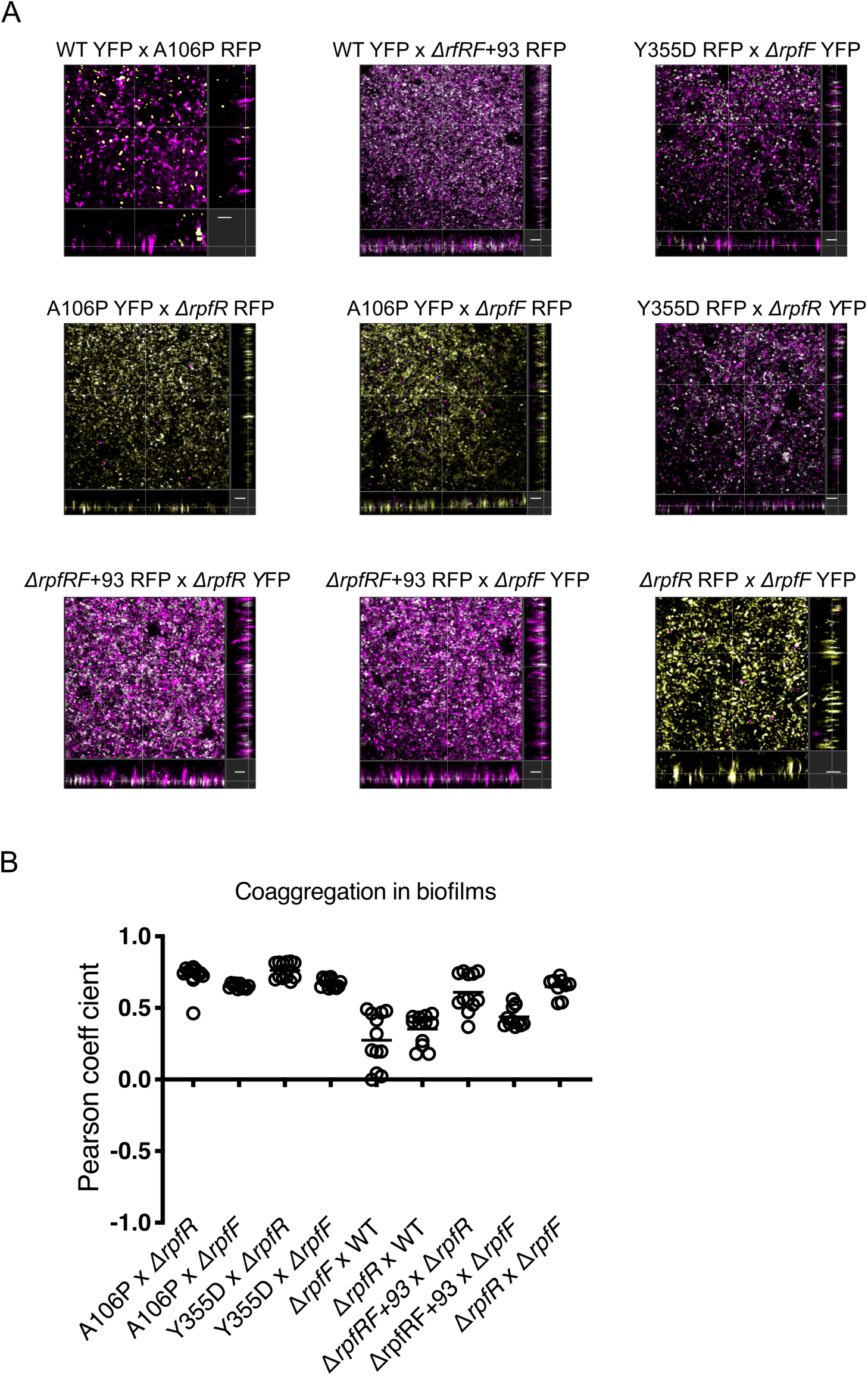
Microscopic biofilm structures and extent of coaggregation. (A) WT and mutants have characteristic structural differences: WT shows isolated small clusters while A106P and Y355D both exhibit large aggregates in biofilm. Note that both *ΔrpfR* and *ΔrpfF* form small clusters amidst larger aggregates produced by A106P and Y355D, implying the interaction between genotypes and formation of a mixed biofilm structure. The biofilms of *ΔrpfRF*+93 with *ΔrpfR* and *ΔrpfF* show small clusters with uniform coverage demonstrating mixed biofilm. *ΔrpfR* and *ΔrpfF* together form large aggregates. Coaggregation in biofilms is represented as white spots and RFP labeled cells are false-colored in magenta for ease of viewing (scale= 10 μm). (B) Coaggregation in biofilms is determined by the Pearson Coefficient (−1= negative correlation, 0= no correlation and 1= positive correlation), where positive values between 0 and 1 indicate the extent of overlap between two channels.

**Fig. S6.**
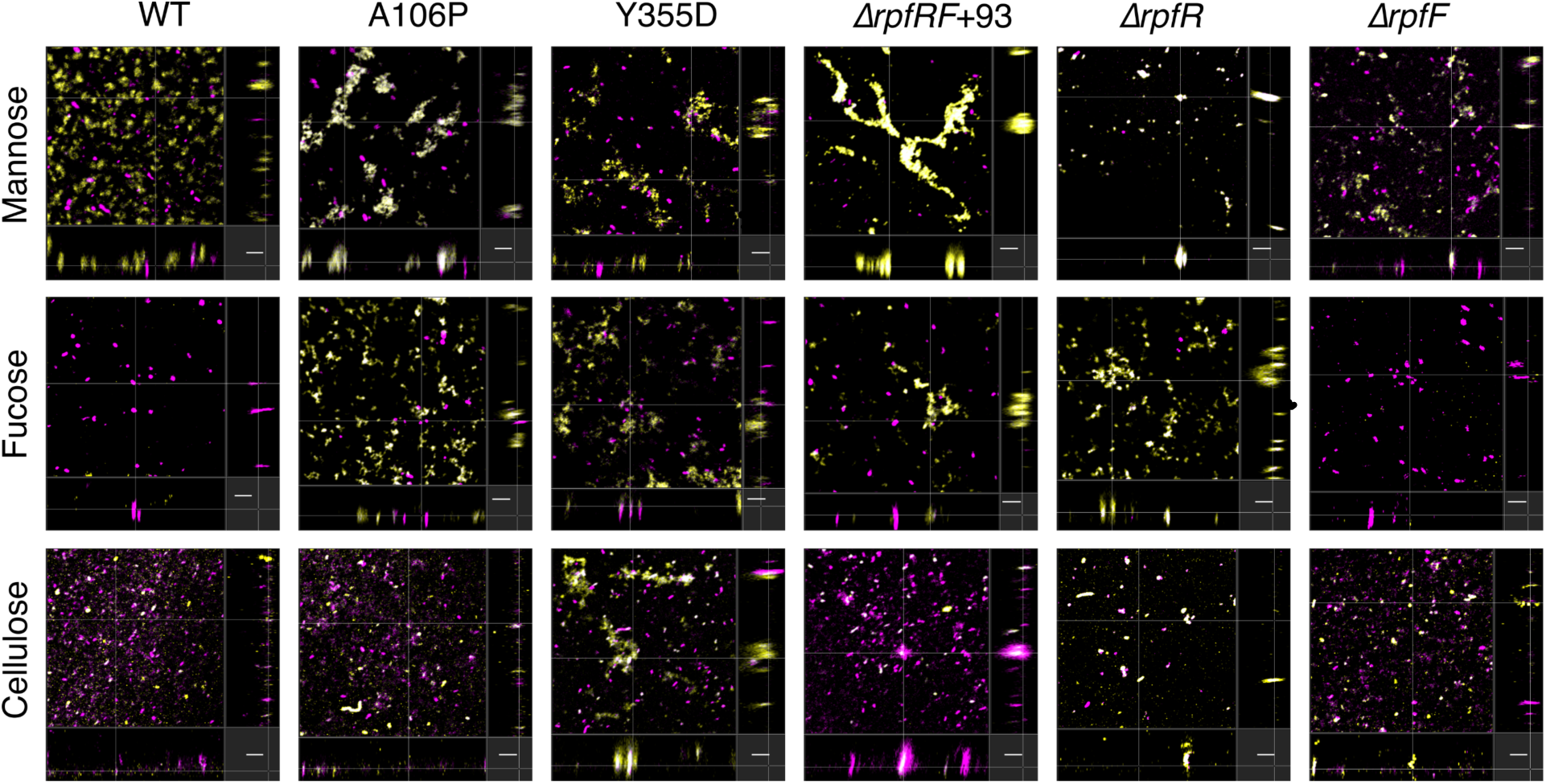
Lectin-based labeling of matrix polysaccharides in biofilms. Fluorescently tagged lectins bind mannose and fucose, and calcofluor white stains cellulose in biofilms. Labels are shown in yellow, while RFP-labeled cells are false-colored in magenta (scale= 10 μm).

**Fig. S7.**
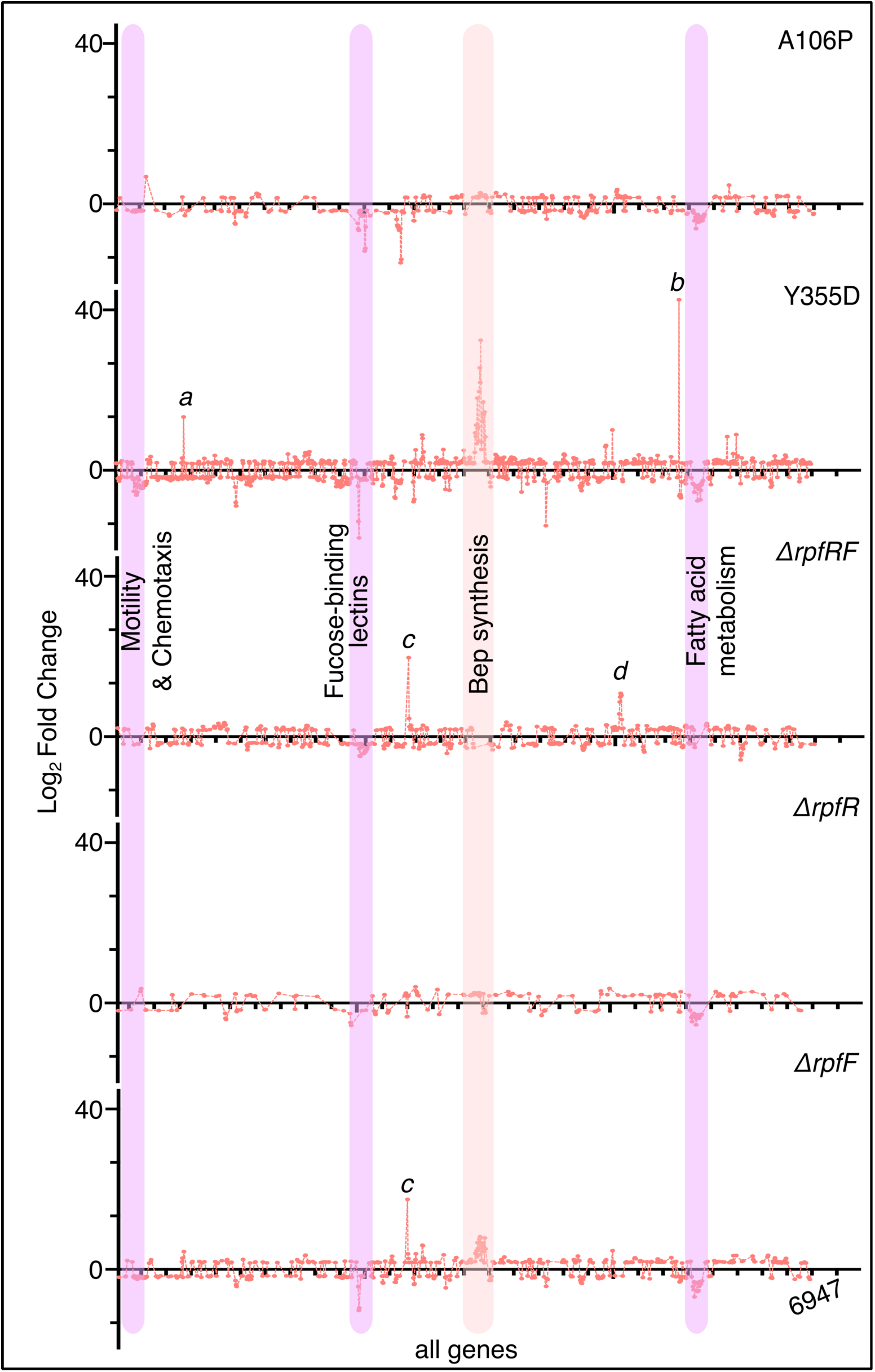
Spectral representation of the mean fold changes in expression. Mean fold-changes in gene expression compared to wild type are calculated from the three biological replicates of each mutant (q value < 0.05). The highlighted (purple: downregulated, orange: upregu lated) gene clusters display parallel shift in 4 or more mutants (highest in *rpfR* Y355D). Specific genes showing significant upregulation or downregulation are marked with letters: a: outer membrane autotransporter, b: Flp/Fap pilin component, c: Bcen2424_3556 (gene adjacent to *rpfF*, function unknown) and d: GTP cyclohydrolase. The raw data is submitted to NCBI BIoproject (Accession number: PRJNA607303).

